# Physoxia influences global and gene-specific methylation in pluripotent stem cells

**DOI:** 10.1101/2022.03.21.484908

**Authors:** Fatma Dogan, Rakad M Kh Aljumaily, Mark Kitchen, Nicholas R. Forsyth

**Affiliations:** The Guy Hilton Research Laboratories, School of Pharmacy and Bioengineering, Faculty of Medicine and Health Sciences, Keele University, Stoke on Trent, UK; Department of Biology, College of Science, University of Baghdad, Baghdad, Iraq

## Abstract

Pluripotent stem cells (PSC) possess unlimited proliferation, self-renewal, and a differentiation capacity spanning all germ layers. Appropriate culture conditions are important for the maintenance of self-renewal, pluripotency, proliferation, differentiation, and epigenetic states. Oxygen concentrations vary across different human tissues depending on precise cell location and proximity to vascularisation. The bulk of PSC culture-based research is performed in a physiologically hyperoxic, air oxygen (21% O_2_) environment, with numerous reports now detailing the impact of physiologic normoxia (physoxia) low oxygen culture in the maintenance of stemness, survival, morphology, proliferation, differentiation potential, and epigenetic profiles. Epigenetic mechanisms affect multiple cellular characteristics including gene expression during development and cell fate determination in differentiated cells. We hypothesized that epigenetic marks are responsive to a reduced oxygen microenvironment in PSCs and their differentiation progeny.

Here, we evaluated the role of physoxia in PSC culture, the regulation of DNA methylation (5mC and 5hmC), and expression of regulatory enzymes DNMTs and TETs. Physoxia enhanced the functional profile of PSC including proliferation, metabolic activity, and stemness attributes. PSCs cultured in physoxia revealed significant downregulation of DNMT3B, DNMT3L, TET1, and TET3 vs. air oxygen, accompanied by significantly reduced 5mC and 5hmC levels. Downregulation of DNMT3B was associated with an increase in its promoter methylation. Coupled to above we also noted decreased HIF1A but increased HIF2A expression in physoxia cultured PSCs, versus air oxygen. In conclusion, PSCs display oxygen-sensitive methylation patterns that correlate with transcriptional and translational regulation of the de novo methylase DNMT3B.

## Introduction

Epigenetics play a key role in pluripotency determination in human pluripotent stem cells (hPSCs) ^1^. A powerful laboratory tool, hPSC are exploited as an in vitro model system to explore differentiation and epigenetic changes during development processes ^2,3^. Human-induced pluripotent stem cells (hiPSCs) are an effective complementary aid for hESCs (embryonic stem cells) studies due to their matched properties including self-renewal and differentiation into multiple cell lineages^4,5^. Further, both hESCs and hiPSCs have a key role to play as cell sources for regenerative medicine and drug screening research ^6,7^. hPSCs display unique epigenetic patterning when compared to differentiated and somatic cells that may contribute to unravelling developmental processes reliant on appropriate gene expression ^8,9^.

Epigenetic mechanisms consist of DNA methylation, histone modifications, and noncoding RNAs. These can modify chromatin and genomic structure and affect gene expression without changing DNA sequence ^10,11^. DNA methyltransferase enzymes (DNMTs) include DNMT1, DNMT3A, DNMT3B, and accessory protein DNMT3L, and establish and maintain DNA methylation patterns in mammals ^12^. The structure, mechanism, and function of DNMTs are well defined. DNMT1 is predominantly involved in the maintenance of DNA methylation during cell division while DNMT3A and DNMT3B, which work in coordination with DNMT1, are responsible for de novo methylation, typically during early development in embryonic stem cells ^13,14^. DNMT3A and DNMT3B expression is high in undifferentiated ESCs, down-regulated post-differentiation, and remains low in adult somatic tissues ^15,16^. DNMT3A and DNMT3B knockout cells can differentiate into the three germ layers (ectoderm, endoderm, mesoderm) ^16^. Further, early-passage DNMT3A and DNMT3B knockout mouse ESCs can initiate and proceed to terminally differentiated cardiomyocytes and hematopoietic cells where DNMT1 is essential for maintenance of differentiation potential ^17,18^. Knockout of DNMT3A and DNMT3B in ESCs does not impact replication potential but does so in differentiated progeny ^18^.

Oxygen is fundamental for cellular processes. hESCs are derived from preimplantation blastocysts that would undergo routine physiological exposure to a 2-5% ranged oxygen environment ^19^ but paradoxically air oxygen remains the predominant culture condition. Physiological oxygen tensions promote maintenance of key properties on embryonic and mesenchymal stem cells, are an essential component of the stem cell microenvironment, and act as a signalling molecule in the regulation of stem cell development ^20^. Physoxia promotes hESC expansion, clonogenicity, transcriptional (mRNA and miRNA) changes, translational changes, genomic stability, altered metabolic activity, migration, and differentiation features ^21–25^.

Methylation of genomic CpG islands is correlated to transcriptional silencing and is required for normal development and hESCs differentiation ^26^. Key transcription factors (Nanog and Oct4/Pou5f1), which maintain stem cell features in ESCs, are unmethylated in undifferentiated stem cells, and become methylated as differentiation progresses ^27^. Physiological oxygen (1%) promote significant reduction of global DNA methylation levels in human colorectal, melanoma, and neuroblastoma cancer cell lines ^28,29^. Moreover, in neuroblastoma this is accompanied by increased TET1 transcription and elevated global 5hmC levels with an accumulation of 5hmC density at hypoxia response genes ^29^. Similarly, hMSC (human mesenchymal stem cell) isolation and continuous exposure to 2% O_2_ resulted in a significant decrease in global 5mC and 5hmC levels, accompanied by reduced expression of DNMT3B and HIF1A but increased expression of HIF2A ^30^. TET1 and TET2 expression is reported as having a key role in cell fate and differentiation potential in undifferentiated ESCs^31^. TET1 activity, for example, is regulated by physiological O_2_ in embryogenesis and is specifically inhibited by low (1%) O_2_ ^32^. Downregulation of TET1 is associated with decreased global 5hmC, both in undifferentiated and 3 days differentiated mESC in 1% O_2_, in addition, silencing TET1 increased expression of pluripotency markers and inhibited mESCs differentiation potential ^32^. Our previous observations detailed reductions in global 5mC, 5hmC levels and TET1 expression in hMSCs cultured in physoxia ^30^. DNMT1 and DNMT3B promoters both possess HIF1A binding sites, suggesting that oxygen levels can play a role in the regulation of DNMT1 and DNMT3B expression via HIF1A transcription factor ^33,34^. Prolonged exposure to low oxygen culture is reported to be associated with decreased HIF1A expression in hESCs ^19^. Consistent with this report decreased DNMT3B and HIF1A expression has been observed after long term culture in 2% O_2_ in hMSCs ^30^. The effects of physoxia on DNA methylation in hPSCs and differentiated progeny have received little attention so far.

We report a significant decrease in global methylation (5mC and 5hmC) and DNMT3B/ TET1 expression consistent with increased promoter-specific methylation patterns linked to physoxia culture in PSCs. hPSCs have great potential to provide new research tools for clinical applications such as drug screening and cell replacement therapies. Understanding the methylation patterning associated with hPSCs differentiation in physiological settings is essential for the development of functionally relevant models.

## Materials and Methods

### Cell culture

SHEF1 and SHEF2 were obtained from the laboratory stock (Guy Hilton Research Centre) and used under approval from the UK Stem Cell Bank (UKSCB) ^35^. Human-induced pluripotent stem cell line (iPSC) (ZK2012L) derived from human dermal fibroblasts was kindly provided by Professor Susan Kimber and Dr Zoher Kapacee, Faculty of Biology Medicine and Health, University of Manchester. PSCs were routinely cultured in E8 medium (Life Technologies) with Essential 8 Supplement (1X) in culture vessels coated with 5mg/ml vitronectin (Recombinant Human Protein; Life Technologies, London, UK). Spontaneous differentiation medium was Knockout DMEM, 10% FBS, 1% NEAA, 1% L-glutamine and β-mercaptoethanol. hPSCs were grown in vitronectin-coated culture plates for 48 hours in E8 medium before switching into a spontaneous differentiation medium. Confluent cells were passaged enzymatically using 0.5mM EDTA (Fisher Scientific, Loughborough, UK) and incubated for 3-5 minutes at 37°C until cells began to detach. Then, cells were collected in pre-warmed media and centrifuged at 1000 rpm for 3 minutes. Next, the supernatant was removed, the pellet re-suspended in fresh media, and seeded at a minimum split ratio of 1:2. Cells were maintained in three different oxygen settings; 21% air oxygen (21% AO), a fully defined 2% O_2_ environment (workstation) (2% WKS), and a standard 2% O_2_ incubator (2% PG) where samples were handled in a standard class II biological safety laminar flow cabinet. Media utilised in either 2% O_2_ setting was deoxygenated to a 2% O_2_ level prior to use using defined Hypoxycool (Baker Ruskinn, Bridgend, UK) cycle settings.

### Metabolic activity of hPSCs

PSCs were seeded into 96-well plates at a density of 1 × 10^4^ cells/cm^2^, incubated for 24 hrs, MTT stock solution (15 μl) added to 150 μl of culture media and further incubated for 4 hours at 37°C. Following incubation, a 125µl aliquot was removed and 50μl of Dimethyl sulfoxide (DMSO) added, then incubated for 45 minutes at 37°C. For Alamar blue assay (Fisher Scientific, Loughborough, UK), 1 × 10^4^ cells/cm^2^ cells were cultured in flat-bottom 96-well plates for 48 hrs. Media was then removed and replaced with 100μl fresh E8 medium supplemented with 10μl Alamar blue reagent and incubated for 4 hours at 37°C. After final incubation steps, both assays were measured with a plate reader (BioTek, Synergy 2) at a 570 nm wavelength. Data are presented as a mean ± standard deviation (SD) of three triplicate experiments.

### Flow Cytometry

Cells were characterised using hPSC imaging kit (Human Pluripotent Stem Cell Marker Antibody Panel kit, SC008, R&D system, UK) with analysis performed on a flow cytometer (Beckman Coulter Cytomics FC 500). hPSCs were labelled with pluripotency markers (SSEA-4, TRA-1-60 and SSEA-1) and 5 × 10^5^ cells used for each experiment. After harvest, pellets were re-suspended in flow cytometry buffer (PBS with 0.5% (w/v) BSA and 2mM EDTA, (Fisher Scientific). The percentage of positive events was determined via gate exclusion of 99% of control events. Flowing Software was used for data analysis.

### DNA methylation analysis

DNeasy Blood and Tissue kit (Qiagen, Manchester, UK) was used to extract genomic DNA isolation from hPSCs. Total 5mC levels within genomic DNA were measured with the MethylFlashTM Methylated DNA Quantification Kit (Epigentek, USA) using 100ng of input DNA. Total 5hmC quantification was performed with the MethylFlashTM Hydroxymethylated DNA Quantification Kit (Epigentek, USA) using 200ng of input DNA. Methylation levels for both assays were established by reading absorbance at A450 nm via a microplate reader (BioTek, Synergy 2) using program Gen5 1.10 for quantification via provided standards.

### Pyrosequencing

Bisulphite conversion of 500 ng input genomic DNA per sample was performed using EZ DNA Methylation-Gold™ Kit (Zymo Research, Orange, CA, USA). Primers specific for DNMT1, DNMT3A, DNMT3B, DNMT3L, TET1, TET2, and TET3 were designed via the PyroMark Q24 Software 2.0. Primer sequences and product sizes are listed in ***Supplemental Information 2*** and supplied by Biomers (Germany). Converted DNA (2-4μl) was used as a template in PCR reactions and amplifications were performed with the GoTaq^®^G2 Flexi DNA Polymerase kit (Promega, Southampton, UK). Cycling parameters were one cycle of 95°C for 5 minutes for initial denaturation followed by touch-down cycling for the first 14 cycles, where the temperature was reduced by 0.5°C in each successive cycle. This was followed by 35 cycles of 95°C for 45 seconds, annealing at 55–63°C for 45 seconds, elongation at 72°C for 30 seconds, and a final elongation step at 72°C for 5 minutes. PCR amplification product quality was confirmed via 2% agarose gel electrophoresis. Biotin-labelled PCR amplicons were captured by mixing PCR products with streptavidin-sepharose beads (GE Healthcare). Vacuum Prep Tool with filter probes was used to capture sepharose beads with biotin labelled PCR amplicons. Filter probes were washed with 70% ethanol solution, then denaturation solution (NaOH) to denature PCR product and washing buffer to purify final biotinylated strands. The filter probes were carefully placed into the Q24 pyrosequencing plate wells and the beads released into an annealing mix. Then, pyroMark Gold Q24 Reagents including the four nucleotides, the substrate and enzyme mix were loaded into a pyrosequencing dispensation cartridge. Pyrosequencing was started after the cartridge and Q24 pyrosequencing plate were inserted into the pyrosequencing instrument. Data were analysed using PyroMark Q24 Software 2.0.

### Gene expression

RNA was isolated from hPSCs with the RNeasy^®^ Mini Kit (Qiagen, Manchester, UK) and concentration quantified using a NanoDropTM 2000/2000c Spectrophotometer (Thermo Scientific). RT-PCR was performed using the QuantiFast SYBR Green OneStep RT-PCR kit (Qiagen, Manchester, UK). PCR primers were ordered from Invitrogen Ltd and ThermoFisher-Scientific (London, UK). Primer sequences and product sizes are listed in ***Supplemental Information 1***. PCR amplification was performed on 25ng of isolated RNA and relative quantification of gene expression was measured using the 2^-ΔΔCT^ method. All primers had 55°C annealing temperature except DNMT3A at 56°C.

### Protein analysis

BCA protein assay (Sigma-Aldrich) was performed to measure protein concentrations. 30μg of total protein was used for western blot analysis using antibodies against DNMT3B (R&D System/ MAB7646, Secondary Anti-mouse IgG-HRP, Cell Signalling/70765), TET1 (ThermoFisher/ GT1462, Secondary Anti-mouse IgG-HRP, Cell Signalling/70765, London, UK), HIF1A (Novusbio/NB100-479, Secondary Anti-rabbit IgG-HRP Cell Signalling/70745, London, UK), HIF2A (Novusbio/NB100-122, Secondary Anti-rabbit IgG-HRP Cell Signalling/70745, London, UK) and GAPDH (Merck/MAB374, Secondary Anti-mouse IgG-HRP, Cell Signalling/70765, London, UK). Blots were imaged using a FluorChem M Imager system.

### Statistical analysis

The data were analysed using statistical software SPSS (IBM SPSS Statistics 21). A one-way analysis of variance (ANOVA) was performed to assess the comparison among the three groups. The threshold for statistical significance was accepted as p<0.05. GraphPad Prism 5 (GraphPad Software, La Jolla, USA) was used for graphical displaying of data. Experimental data are represented as mean ± standard deviation (SD) and describe a minimum of 3 independent replicates.

## Results

### Reduced oxygen increases proliferation and metabolic activity in hPSC

We determined the proliferation of hPSCs cultured in air oxygen (21% AO) and reduced oxygen conditions (2% PG and 2% WKS) over a 6-day period. A significant increase in proliferation between days 2 and 6 in physoxia settings was noted comparison to those cultured in air oxygen (AO, p<0.05). Metabolic activity was explored via MTT and Alamar blue assays. A significant increase was again noted for hPSC cultured in physoxia when compared to AO (p<0.05). SHEF1 and ZK2012L displayed a significant increase in MTT after day 3 while SHEF2 cells displayed a significant increase in MTT after day 5. Alamar blue indicated increased metabolic activity and proliferation of cells cultured under both reduced oxygen conditions from day 4 (***<u>Figure S1</u>***).

### Increased expression of TRA-160 in physoxia cultured hPSCs

The expression of SSEA-4, TRA-1-60 (pluripotency markers) and SSEA-1 (differentiation marker) were then explored via flow cytometry in undifferentiated hPSCs. As we expected, hPSCs displayed low levesl of SSEA-1 expression coupled to high levels of SSEA-4 and TRA-1-60. No significant difference between conditions for SSEA-4 and SSEA-1 were observed. However, we noted significantly higher expression of TRA-1-60 in SHEF1 cells cultured in 2% PG (88.41% ± 1.79, p>0.01), SHEF2 cells in 2% WKS (85.64% ± 1.26, p>0.01) and ZK2012L cells in 2% PG, 2% WKS (76.84% ± 10.84, p>0.01 and 69.61% ± 9.77, p>0.05) versus 21% AO (67.23% ± 10.57, 62.71% ± 18.18 and 53.48% ± 7.66) (***Fig 1***).

**Figure 1.**
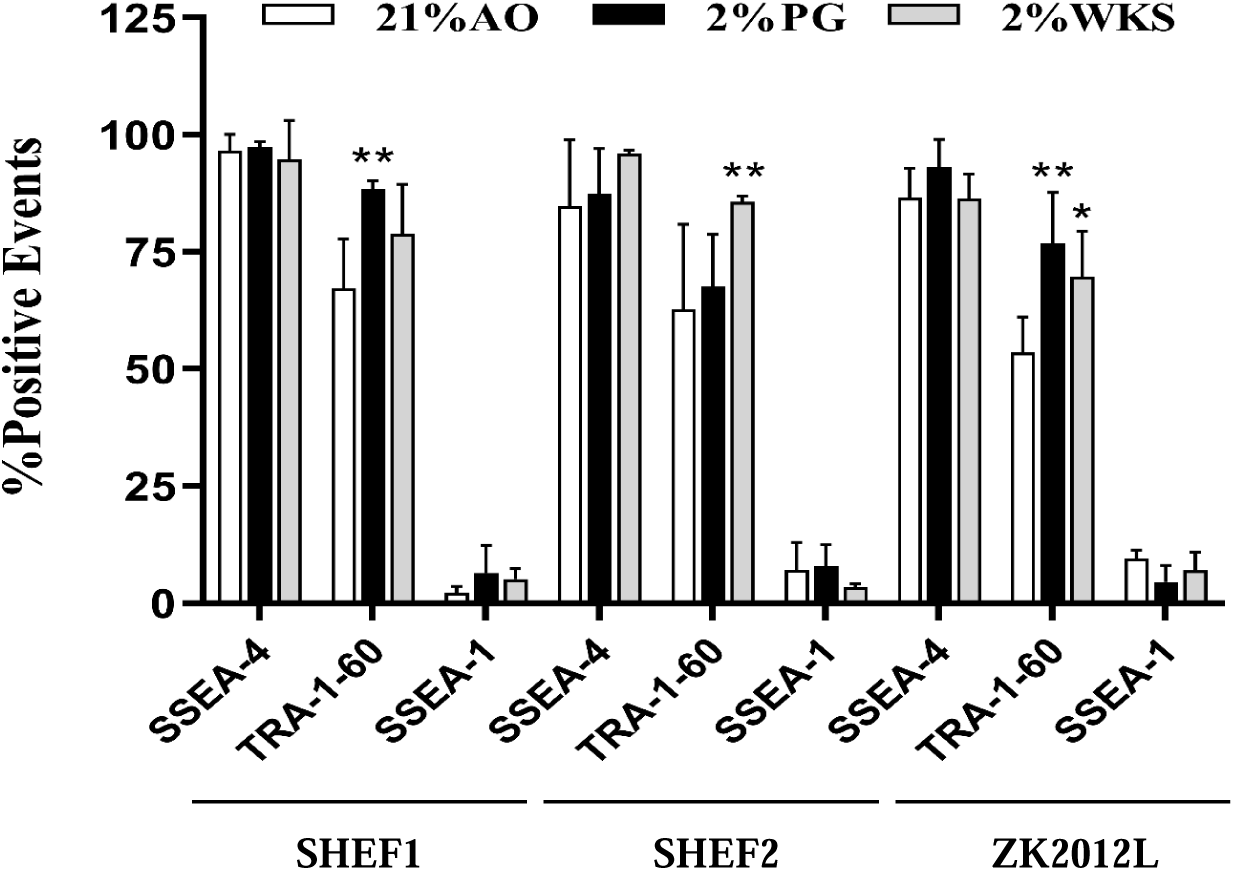
Immunophenotypic characterisation of PSCs. Flow cytometry-based evaluation of expression of pluripotency markers. All PSCs were analysed for SSEA-4, TRA-1-60, SSEA-1 expression in both reduced oxygen (2% PG and 2% WKS) and 21% AO conditions. Y-axis indicates sample names and surface markers. The X-axis shows percentage of positive events. Data are presented as a mean (n=3), *P<0.05, **p<0.01 vs 21% AO, and error bar indicates standard deviation (SD).

### Elevated OCT-4 and SOX-2 expression in physoxia cultured hPSCs

Expression of pluripotency factors OCT-4, NANOG and SOX2 was assessed transcriptionally in different oxygen conditions (21% AO, 2% PG and 2% WKS). OCT-4 expression was increased in 2% PG and 2% WKS in undifferentiated SHEF1 (1.30 ± 0.23 and 1.76 ± 0.50, respectively), SHEF2 (1.10 ± 0.23 and 1.51 ± 0.10, respectively), and ZK2012L (1.84 ± 0.20 and 1.64 ± 0.21, p>0.01, respectively) compared with AO. During differentiation, OCT-4 expression decreased in differentiated SHEF1 and SHEF2 at days 5, 10, and 20 in physoxia. SHEF1 and SHEF2 differentiated in 2% PG showed significant decreases at day 20 (0.33 ± 0.28 and 0. 0.30 ± 0.16, p>0.01, respectively) and SHEF2 in 2% WKS at day 5 and 20 (0.66 ± 0.11, p>0.05 and 0.38 ± 0.19, p>0.01) displayed a significant reduction compared to 21% AO. Undifferentiated ZK2012L cells showed increased OCT-4 expression in 2% PG and 2% WKS (1.83 ± 0.20 and 1.64 ± 0.21, p>0.01). Pooled hPSCs data indicated a significant increase in OCT-4 expression in undifferentiated cells in 2% WKS (1.64 ± 0.13, p>0.05) in comparison to AO with decreased expression at differentiation days 5, 10, and 20. NANOG expression decreased in PSCs cultured in reduced oxygen environments in undifferentiated and 5 days differentiated SHEF1, SHEF2, and ZK2012L versus AO, significantly in undifferentiated SHEF1 in 2% PG (0.51 ± 0.31, p>0.05). Pooled data indicated that NANOG expression was significantly lower in undifferentiated cells cultured in 2% PG and 2% WKS (0.57 ± 0.1 and 0.64 ± 0.11, p>0.01, respectively) and after 5 days differentiation in 2% PG (0.63 ± 0.09, p>0.05) compared to AO. An overall increased SOX-2 expression was noted in PSCs cultured in reduced oxygen conditions. There was a significant increase in undifferentiated SHEF1 and SHEF2 cells cultured in 2% WKS (2.08 ± 0.62 and 1.55 ± 0.19, p>0.05). Also, 10 days differentiated ZK2012L displayed a significant elevation in 2% WKS (2.98 ± 1.01, p>0.05) versus AO. Pooled hPSC data indicated significantly elevated SOX-2 expression in undifferentiated cells cultured in 2% WKS environment (1.96 ± 0.36, p>0.01) in comparison to AO (Fig ***2***).

**Figure 2.**
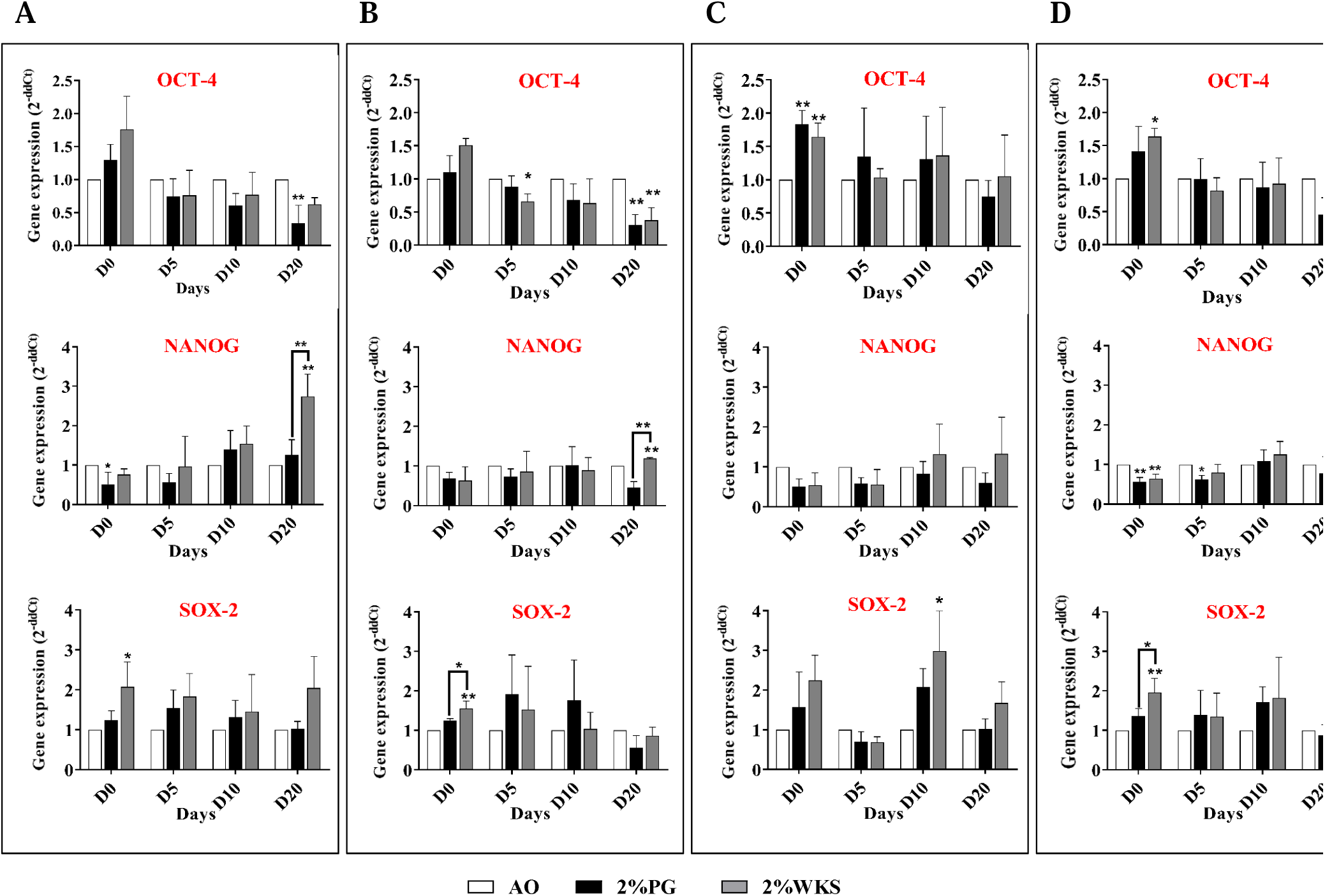
QRT-PCR analysis of OCT-4, SOX-2 and NANOG expression. **(A)** SHEF1, **(B)** SHEF2, **(C)** ZK2012L, and **(D)** Pooled PSCs-profile, (SHEF1, SHEF2 and ZK2012L). The internal control, ACTB, was used to normalize expression. Y-axis shows the relative changes in *2*^-ΔΔ*CT*^ of air oxygen (AO) to physoxia cultured cells. The X-axis indicates time (days). Data are presented as a mean (n=3), *P<0.05, **p<0.01 vs 21% AO, connecting lines indicate significance between conditions, error bars indicate SD.

### Physoxia decreases global DNA methylation in hPSCs

DNA from undifferentiated and differentiated hPSCs was isolated to explore global DNA methylation levels in 21% AO, 2% PG, and 2% WKS. Decreased global DNA methylation in hPSCs cultured in physoxia (2% PG and 2% WKS) versus 21% AO was noted. Further, global DNA methylation was reduced in a time-dependent manner as differentiation progressed in PSCs.

Undifferentiated SHEF1 cultured in either 2% PG (0.82 ± 0.06, p<0.05) or 2% WKS (0.73 ± 0.07, p<0.01) showed significantly decreased 5mC levels versus 21% AO (1.01 ± 0.041). The level of 5mC in day 20 differentiated SHEF1 was significantly decreased in 2% WKS (0.52 ± 0.09, p<0.05) in comparison to 21% AO (0.72 ± 0.074) (***Fig 3***.***A)***. Undifferentiated and day 20 differentiated SHEF2 in 2% PG displayed significantly decreased 5mC (0.74 ± 0.03 and 0.51 ± 0.02, p<0.05) versus 21% AO (0.84 ± 0.05 and 0.70 ± 0.08), respectively. There was a significant decrease in 5mC content at days 10 and 20 in differentiated SHEF2 (0.50 ± 0.03 and 0.41 ± 0.06, p<0.01) in 2% WKS ***(Fig 3***.***B)***.

**Figure 3.**
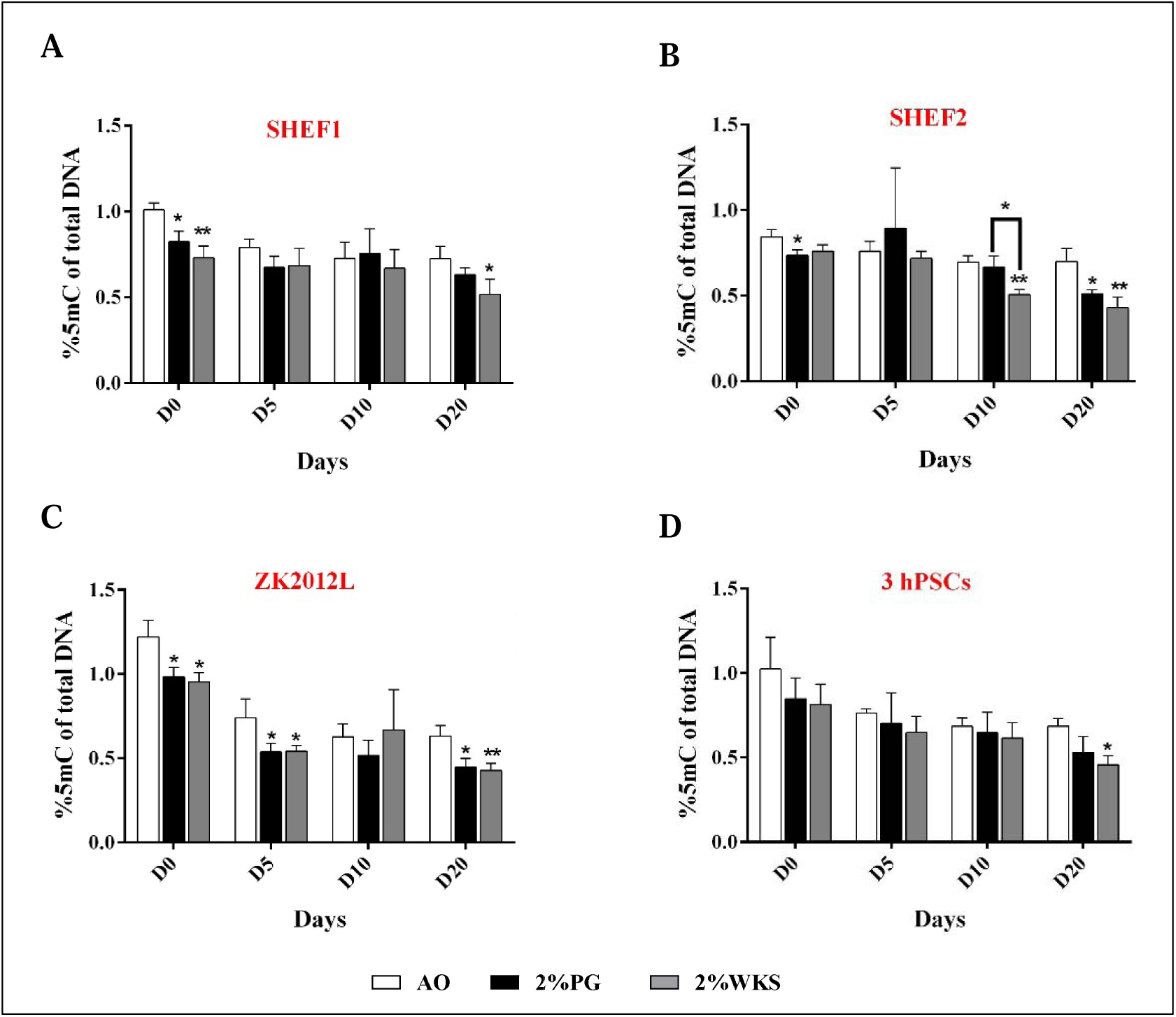
Global 5mC levels in PSCs. **(A)** SHEF1, **(B)** SHEF2, **(C)** ZK2012L, and **(D)** Pooled PSCs-profile, (SHEF1, SHEF2 and ZK2012L). The methylated DNA fragments of PSCs were analysed using MethylFlash Global DNA Methylation (5-mC) Quantification Kit. Y-axis shows the absorbance (450 nm) of 5-methylcytosine. The X-axis indicates different time points (days). Data are presented as a mean (n=3), *P<0.05, **p<0.01 vs 21% AO, connecting lines indicate significance between conditions, error bars indicate SD.

Undifferentiated ZK2012L cultured in 2% PG (0.98 ± 0.06, p<0.05) and 2% WKS (0.95 ± 0.05, p<0.05) showed significantly decreased 5mC levels versus 21% AO (1.22 ± 0.09). Days 5 (0.54 ± 0.05 and 0.54 ± 0.03, p<0.05) and 20 (0.45 ± 0.05 and 0.43 ± 0.04, p<0.05) differentiated cells showed a significant reduction in global 5mC level in 2% PG and 2% WKS, respectively ***(Fig 3***.***C)***. Pooled data from three PSCs (SHEF1, SHEF2 and ZK2012L) indicated reduced global 5mC levels in undifferentiated 2% PG and 2% WKS with a significant decrease in 2% WKS at day 20 (0.46 ± 0.5) in comparison to 21% AO (0.69 ± 0.5) (***Fig 3***.***D***).

Significant decreases were observed in global 5hmC DNA levels in undifferentiated SHEF1 cultured in 2% PG and 2% WKS physoxia environments (0.08 ± 0.006, p<0.05 and 0.06 ± 0.008, p<0.01, respectively) compared to AO (0.13 ± 0.03). Significant decreases at days 10 and 20 differentiation in 2% WKS (0.05 ± 0.01, p<0.05 and 0.04 ± 0.008, p<0.05), respectively, versus 21% AO (0.07± 0.009) were noted ***(Fig 4***.***A)***. Undifferentiated SHEF2 in 2% WKS cells showed a significant decrease (0.14 ± 0.02, p<0.05) in comparison to cells cultured in 21% AO (0.21 ± 0.03). ZK2012L cultured in 2% WKS showed a significant reduction in the percentage of 5hmC at days 0, 5, and 20 (0.13 ± 0.02, 0.08 ± 0.02 and 0.06 ± 0.01, p<0.05, respectively) versus 21% AO (0.21 ± 0.04, 0.12 ± 0.02 and 0.09 ± 0.02) ***(Fig 4***.***B)***. ZK2012L displayed a significant decrease in 2% PG at day 20 (0.06 ± 0.01) in comparison to 21% AO (0.15± 0.02) (***Fig 4***.***C***). Lastly, no significant differences was observed from pooled PSCs data, but an overall reduction in global 5hmC levels in cells cultured under physoxia was noted in undifferentiated and differentiated cells (***Fig 4***.***D)***.

**Figure 4.**
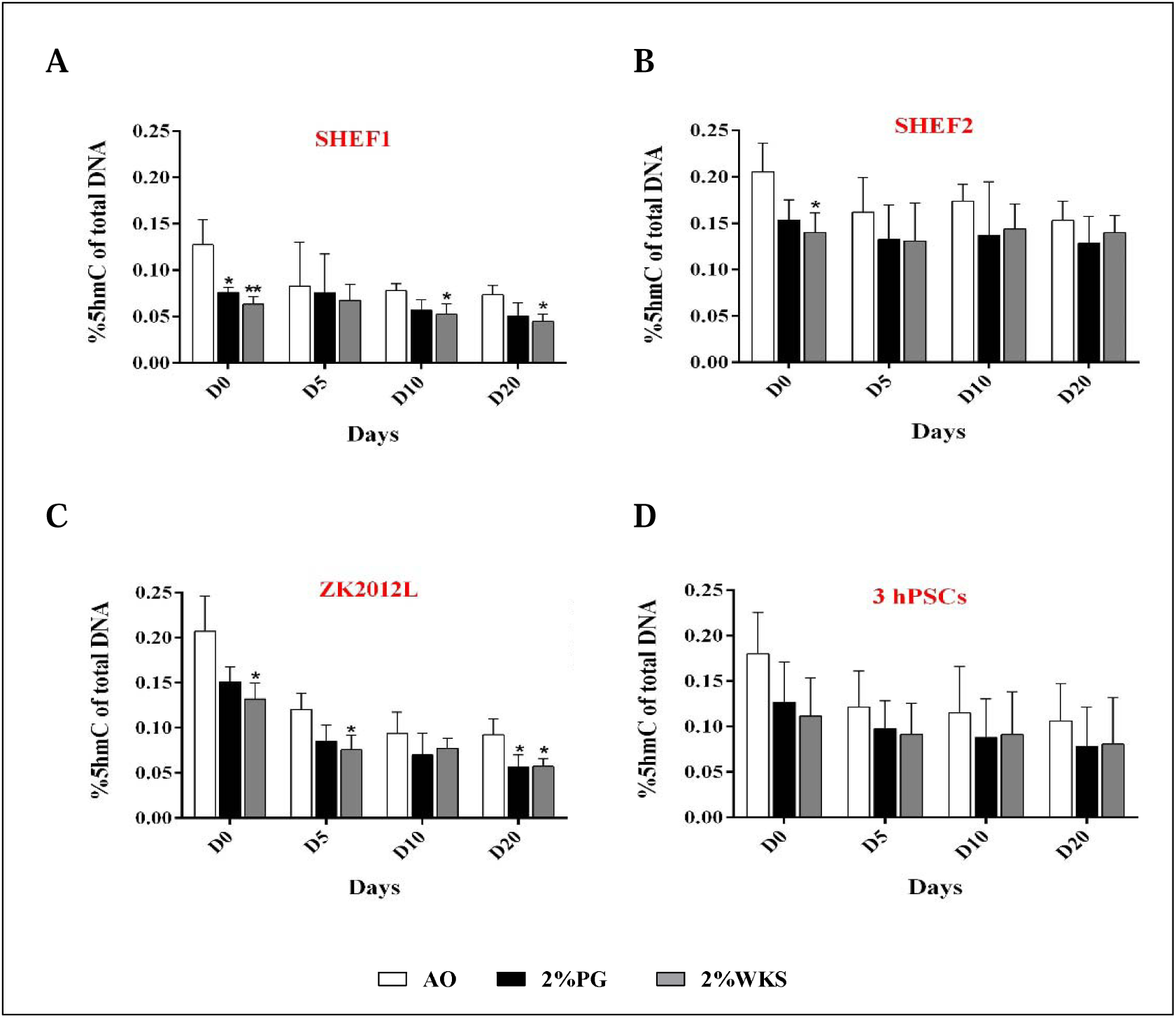
Global 5hmC levels PSCs. **(A)** SHEF1, **(B)** SHEF2, **(C)** ZK2012L, and **(D)** Pooled PSCs-profile, (SHEF1, SHEF2 and ZK2012L). The methylated DNA fragments of PSCs were analysed using MethylFlash Global Hydroxymethylated DNA (5-hmC) Quantification Kit. Y-axis shows absorbance (450 nm) of 5-hydroxymethylcytosine. The X-axis indicates different time points (days). Data are presented as a mean (n=3), *P<0.05, **p<0.01 vs 21% AO, connecting lines indicate significance between conditions, error bars indicate SD.

### Decreased DNMT3B, TET1 and TET3 Gene Expression Associates with Physoxic Culture

No changes in expression of DNMT1 or TET2 were noted during the differentiation of hPSCs. DNMT3A expression was significantly decreased in differentiated SHEF1 at day 5 in 2% WKS (0.65-fold ± 0.22, p<0.05) vs. 21% AO. Undifferentiated SHEF2 and ZK2012L cells displayed reduced DNMT3A gene expression in 2% PG and 2% WKS (0.59-fold ± 0.03 and 0.65-fold ± 0.13, p<0.05, respectively) in comparison to 21% AO.

Decreased DNMT3B expression was noted in undifferentiated SHEF1 in 2% PG and 2% WKS (0.63-fold ± 0.21 and 0.48-fold ± 0.15, p<0.05, respectively), and during spontaneous differentiation at days 5, 10, and 20 in 2% WKS (0.45-fold ± 0.16, p<0.01, 0.50-fold ± 0.19 and 0.49-fold ± 0.21, p<0.05, respectively) versus AO (***Fig 5***.***A)***. Undifferentiated SHEF2 in 2% PG and 2% WKS had reduced DNMT3B expression (0.26-fold ± 0.18 and 0.38-fold ± 0.22, p<0.01, respectively), and after 10 days differentiated (0.52-fold ± 0.19 and 0.36-fold ± 0.24, p<0.05, respectively) in 2% PG and 2% WKS and at day 20 in 2% WKS (0.16-fold ± 0.03, p<0.01) in comparison to those cultured in 21% AO (***Fig 5***.***B)***. Similar reductions were observed in undifferentiated ZK2012L in 2% PG and 2% WKS (0.50-fold ± 0.14 and 0.45-fold ± 0.17, p<0.01, respectively), and after 5 (0.66-fold ± 0.16, p<0.05 and 0.41-fold ± 0.17, p<0.01, respectively) and 20 days (0.55-fold ± 0.17) differentiation versus 21% AO (***Fig 5***.***C)***. Pooled data from all PSCs tested showed a significant reduction in DNMT3B expression in undifferentiated hPSC cultured in either 2% PG or 2% WKS conditions (0.46 ± 0.18 and 0.44 ± 0.05, p<0.01, respectively), and after 20 days differentiation (0.67 ± 0.07, p<0.05 and 0.40 ± 0.21, p<0.01, respectively) in comparison to 21% AO (***Fig 5***.***D)***.

**Figure 5.**
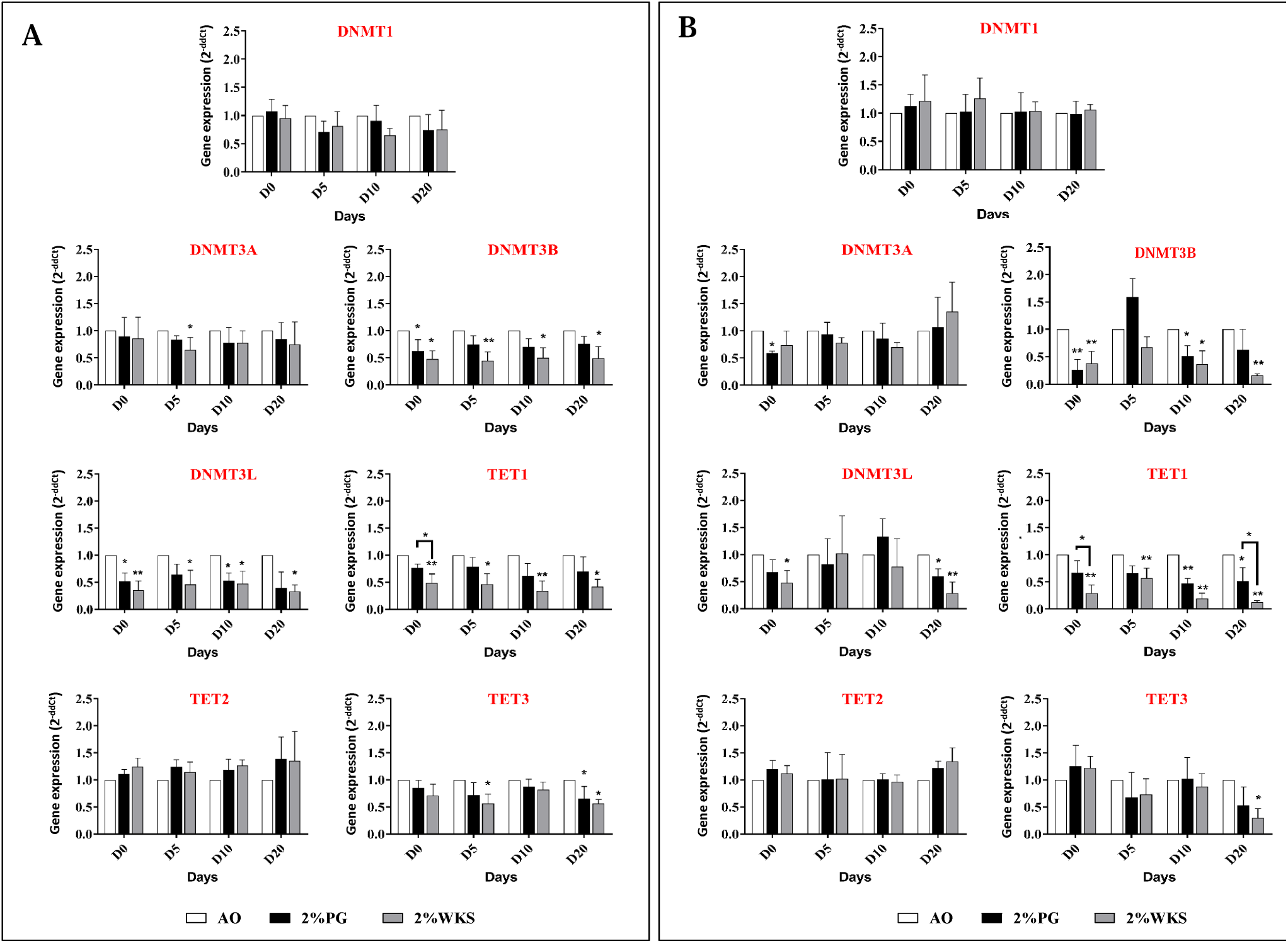
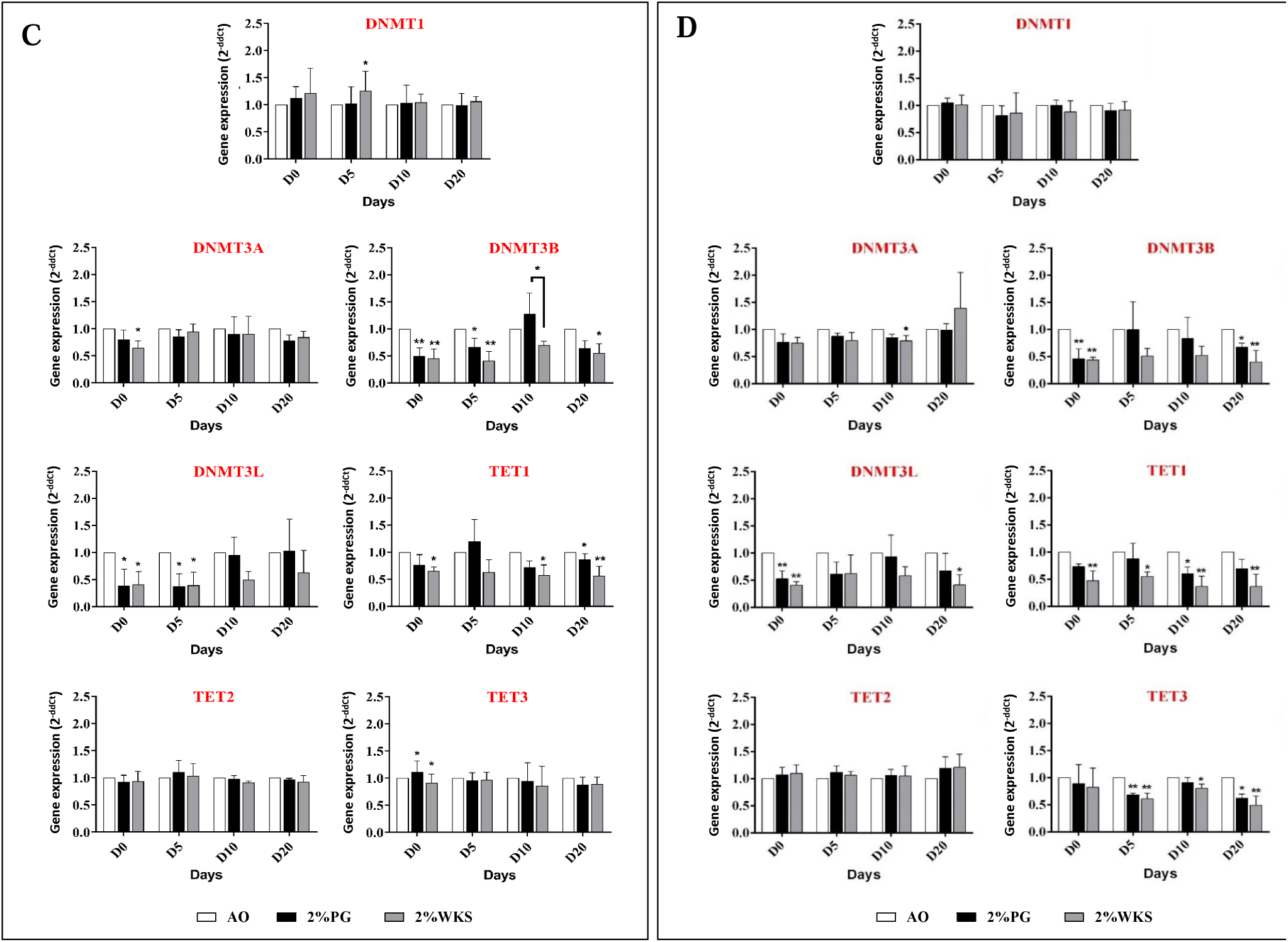
QRT-PCR analysis of DNMTs and TETs in PSCs. Gene expression of DNMTs and TETs was performed **(A)** SHEF1, **(B)** SHEF2, **(C)** ZK2012L, and **(D)** Pooled PSCs lines, (SHEF1, SHEF2 and ZK2012L) in 21% AO and physiological oxygen conditions (2% PG and 2% WKS). The internal control ACTB was used for normalization. Y-axis shows the relative changes in *2*^-ΔΔ*CT*^ of air oxygen to physoxia. The X-axis indicates time (days). Data are presented as a mean (n=3), *P<0.05, **p<0.01 vs 21% AO, connecting lines indicate significance between conditions, error bars indicate SD.

DNMT3L expression was significantly decreased in undifferentiated, and 10 days differentiated SHEF1 in 2% PG (p<0.05) and 2% WKS (p<0.01 and p<0.05, respectively), and in 2%WKS at days 5 and 20 (p<0.05) in comparison to 21% AO (***Fig 5***.***A)***. Decreased expression of DNMT3L was noted in undifferentiated 2% WKS (p<0.05) and 20 days differentiated SHEF2 in 2% PG (p<0.05) and 2% WKS (p<0.01) (***Fig 5***.***B)***. Further, DNMT3L expression was decreased in undifferentiated (p<0.05), and day 5 differentiated ZK2012L cells (p<0.05) in both 2% PG and 2% WKS (***Fig 5***.***C)***. Pooled data from all three PSCs showed a significant reduction (p<0.01) in DNMT3L expression in undifferentiated hPSCs cultured in both 2% PG and 2% WKS and day 20 differentiated cells in 2% WKS (p<0.05) (***Fig 5***.***D)***.

TET1 expression was reduced in undifferentiated SHEF1 (0.49-fold ± 0.17, p<0.01), and after 5 (0.46-fold ± 0.19, p<0.05), 10 (0.34-fold ± 0.18, p<0.01) and 20 days differentiation (0.41-fold ± 0.14, p<0.05) in 2% WKS in comparison to AO (***Fig 5***.***A)***. TET1 expression was reduced in undifferentiated SHEF2 in 2% WKS (0.29-fold ± 0.15, p<0.01), day 5 (0.57-fold ± 0.18, p<0.01), day 10 (0.20-fold ± 0.10, p<0.01), and day 20 (0.13-fold ± 0.03, p<0.01) differentiation, and in 2% PG at day 10 (0.47-fold ± 0.09, p<0.01) and day 20 (0.51-fold ± 0.24, p<0.05) (***Fig 5***.***B)***. TET1 expression in ZK2012L cells was decreased in undifferentiated cells (0.65-fold ± 0.08, p<0.05) and day 10 differentiated cells (0.57-fold ± 0.19, p<0.05) in 2% WKS, and in day 20 with both 2% PG and 2% WKS (0.87-fold ± 0.10, p<0.05 and 0.57-fold ± 0.17, p<0.01, respectively) (***Fig 5***.***C)***. Pooled PSC indicated a significant decrease in TET1 expression in undifferentiated (p<0.01) and differentiated cells at days 5 (p<0.05), 10, and 20 (p<0.01) in 2% WKS and 10 days differentiation in 2% PG (p<0.05) versus AO (***Fig 5***.***D)***.

Finally, TET3 expression was significantly reduced in day 5 differentiated SHEF1 (p<0.05) in 2% WKS and day 20 in 2% PG and 2% WKS (p<0.05) in comparison to those cultured in 21% AO. Significant decreases in day 20 differentiated SHEF2 cultured in 2% WKS (p<0.05) and undifferentiated ZK2012L cells with 2% PG and 2% WKS (p<0.05) were also noted in comparison to AO. Pooled hPSCs data demonstrated decreased TET3 expression at day 5 (p<0.01), day 10 (p<0.05), and day 20 (p<0.01) differentiation in 2% WKS, and both days 5 (p<0.01) and 20 in 2% PG (***Fig 5***).

### Downregulated DNMT3B and TET1 Protein Expression in PSCs

Transcriptional analysis outcomes were confirmed by western blot of undifferentiated and day 20 differentiated hPSCs. A reduction of DNMT3B and TET1 expression in undifferentiated and 20 days differentiated stem cells in physiological oxygen conditions was noted in correlation with transcriptional reductions of DNMT3B and TET1 (***Fig 6)***.

**Figure 6.**
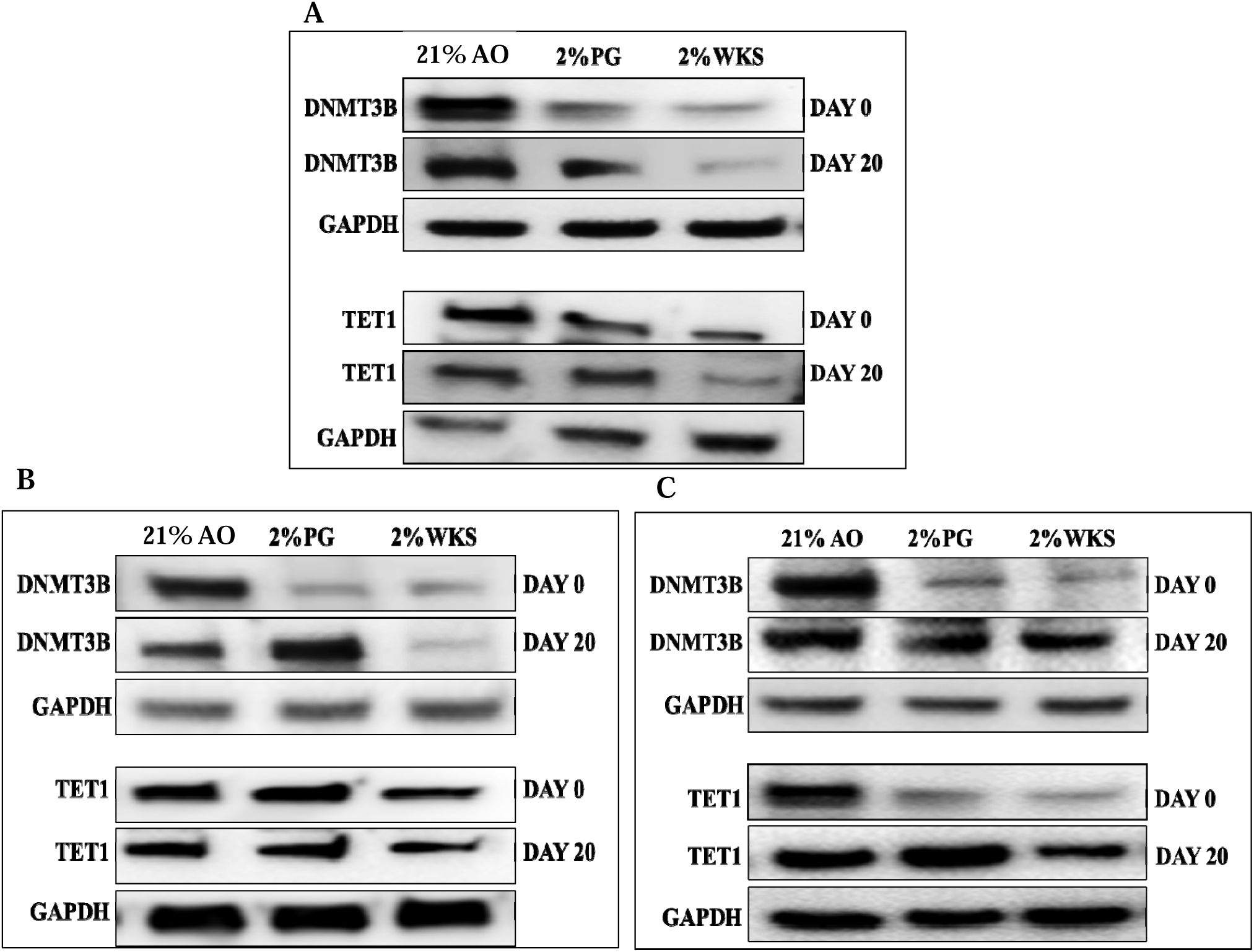
DNMT3B and TET1 protein expression in PSCs. **(A)** SHEF1, **(B)** SHEF2, and **(C)** ZK2012L. Protein samples were isolated from undifferentiated PSCs and 20 days differentiated cells exposed to air oxygen and physoxia conditions. GAPDH was used as a control.

### Physoxia increased DNMT3B AND DNMT3L promoter methylation in hPSCs

The mean level of DNMT3B promoter methylation was higher in undifferentiated SHEF1 in 2% PG (23%, p<0.05) and 2% WKS (32%, p<0.01) versus 21% AO (11%). Methylation levels on DNMT3B promoter were also elevated at differentiation days 5 in 2% PG and 20 in 2% WKS (25%, p<0.05 and 25%, p<0.01) compared to 21% AO (13% and 11%). The percentage of DNMT3L promoter methylation was significantly higher in undifferentiated (86%, p<0.01) and day 10 (90%, p<0.05) and 20 (90%, p<0.01) differentiated cells in 2% WKS compared with 21% AO (70%, 75% and 73%, respectively) (***Fig 7***.***A)***.

**Figure 7.**
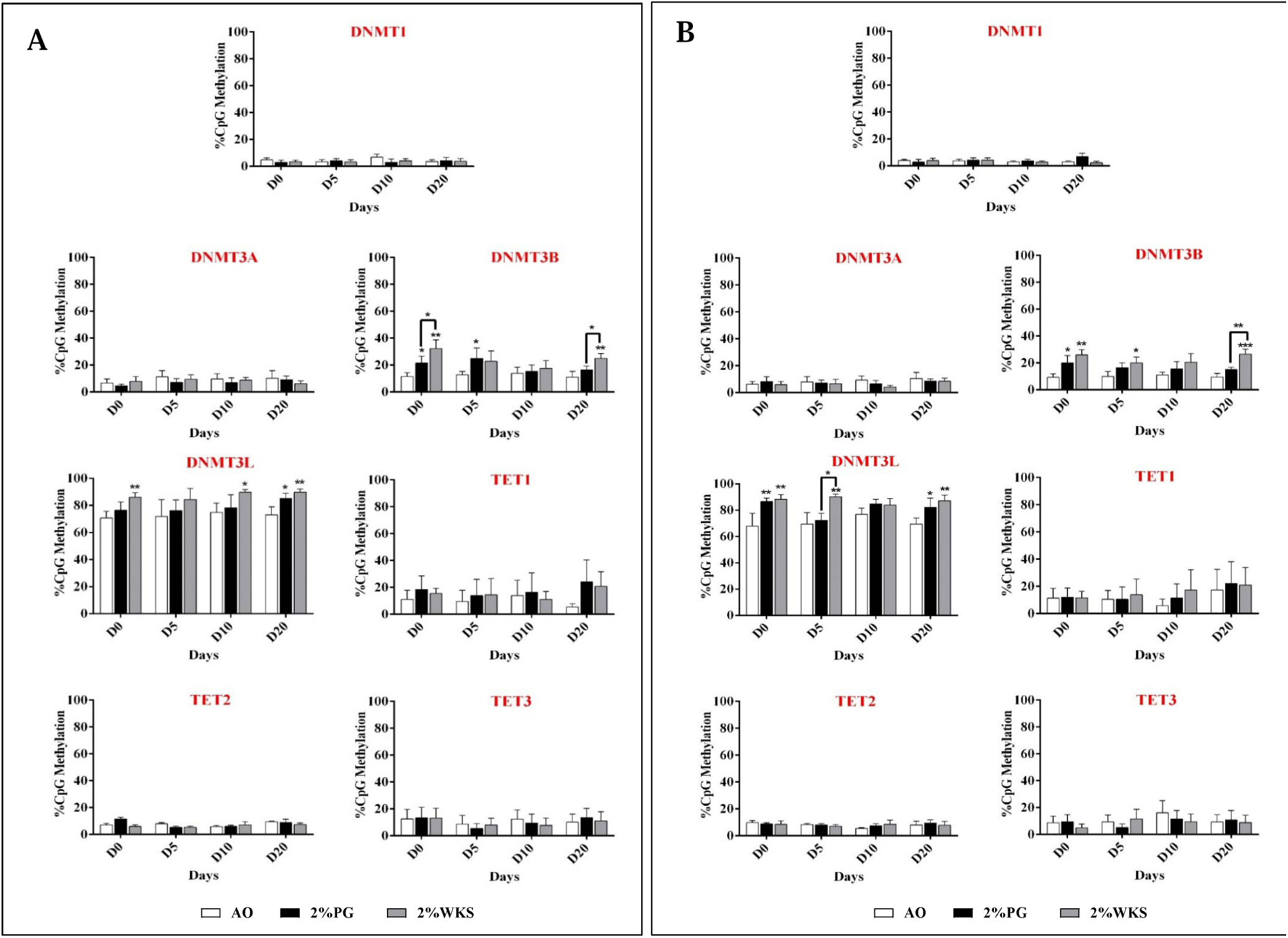
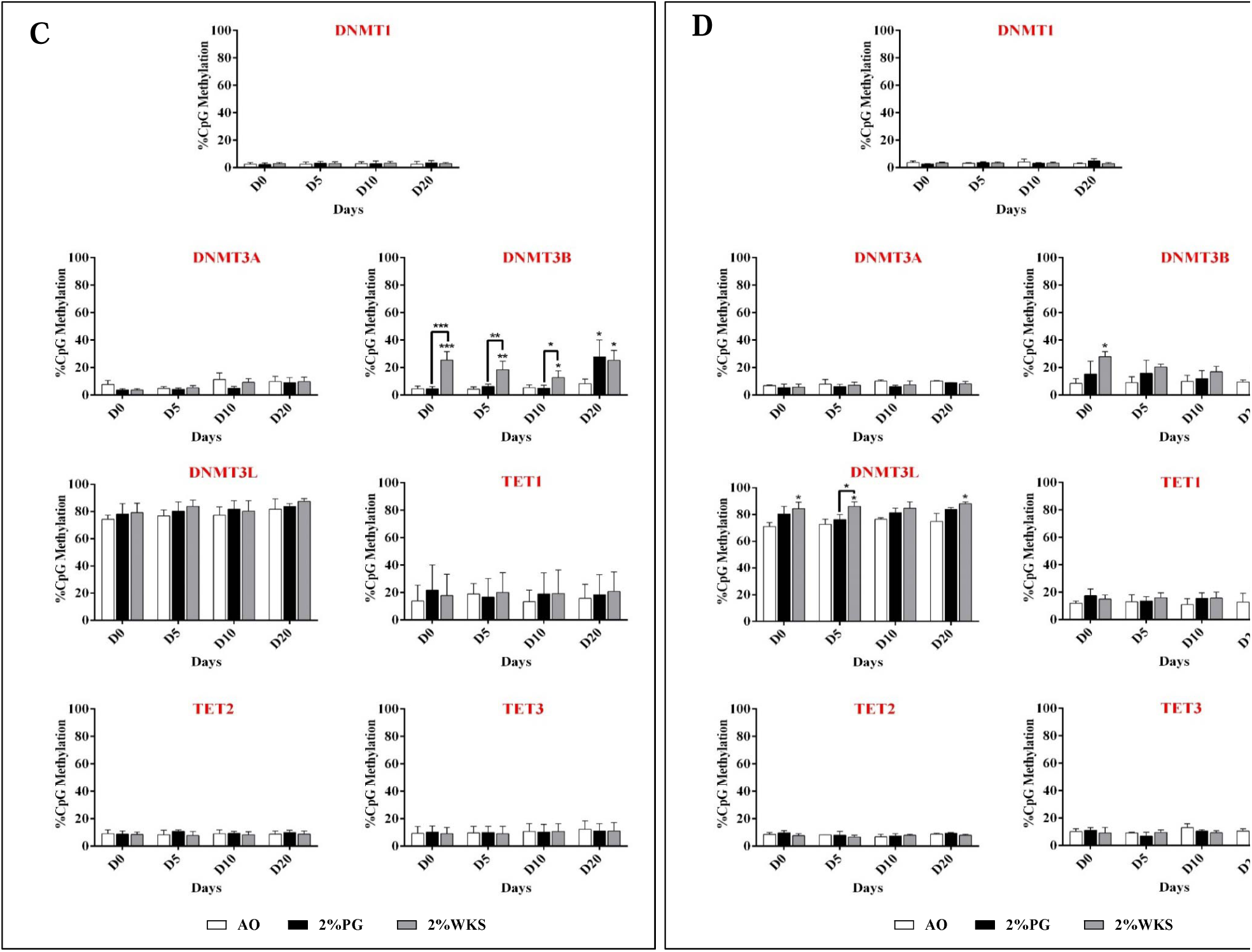
DNMT and TET promoter methylation in PSCs. **(A) SHEF1 and (B) SHEF2, (C)** ZK2012L, and **(D)** Pooled PSCs (SHEF1, SHEF2 and ZK2012L). CpG island methylation in DNMTs and TETs promoters were evaluated using pyrosequencing. Y-axis represents DNA methylation percentage at CpG regions. The X-axis represents time (days). Data are presented as mean (n=3), *P<0.05, **p<0.01 and ***p<0.001 vs 21% AO, connecting lines indicate significance between conditions, error bars indicate SD.

A significant increase in DNMT3B promoter methylation was noted in undifferentiated SHEF2 cells cultured in 2% PG (20%, p<0.05) and 2% WKS (26%, p<0.01) versus 21% AO (9%). In addition, there was a significant elevation in methylation at days 5 (20%, p<0.05) and 20 (27%, p<0.01) in 2% WKS compared with 21% AO (10% and 9%, respectively). Furthermore, DNMT3L showed a significant increase in methylation in 2% PG (87%, p<0.01) and 2% WKS (89%, p<0.01) in undifferentiated SHEF2 cells compared with 21% AO (68%). There was a significant increase in methylation level at day 5 (90%, p<0.01) and 20 in 2%PG and 2%WKS (82%, p<0.05 and 87%, p<0.01, respectively) compared with 21% AO (70%) (***Fig 7***.***B)***.

Undifferentiated ZK2012L (26%, p<0.001) and differentiated cells at day 5 (19%, p<0.01), 10 (13%, p<0.05), and 20 (25%, p<0.05) cultured in 2% WKS had significantly higher DNMT3B promoter methylation levels compared with 21% AO (5%, 4%, 5% and 8%, respectively). No significant change was noted in the methylation of DNMT3L for ZK2012L cells (***Fig 7***.***C)***. Combined data from three PSCs showed that there was a significant increase in undifferentiated (28%, p<0.05) and day 20 differentiated (26%, p<0.01) cells cultured under 2% WKS versus 21% AO (10% and 8%). A significant increase was noted in DNMT3L methylation in undifferentiated cells and in day 5 and 20 from differentiation (85%, 86% and 88%, p<0.05) in 2% WKS conditions in comparison to 21% AO (71%, 73% and 75%) respectively. In general, no significant change was noted in the promoter methylation level of DNMT1, DNMT3A, TET1, TET2 and TET3 genes in either SHEF1, SHEF2, or ZK2012L cells (***Fig 7***.***D)***.

### Physoxia increased HIF2A protein expression in PSCs

Undifferentiated SHEF1 displayed significant downregulation of HIF1A transcript in physoxia (2%PG and 2%WKS, p<0.01), and at days 5 (p<0.01 and p<0.05, respectively), 10 (p<0.05), and 20 of differentiation (p<0.01 and p<0.05, respectively), vs. 21% AO (***Fig 9***.***A)***. A significant decrease was noted in gene expression of HIF1A in undifferentiated SHEF2, and days 5, 10 and 20 differentiated (p<0.05) in 2% PG, and 5, 10 and 20 days differentiated SHEF2 in 2% WKS (p<0.01, p<0.01 and p<0.05, respectively) (***Fig 9***.***B)***. We also noted a significant decrease in undifferentiated ZK2012L (p<0.01, p<0.01), and at differentiation days 5 (p<0.05, p<0.01), 10 (p<0.05, p<0.01) and 20 (p<0.01, p<0.01) in 2% PG and 2% WKS in comparison to 21% AO (***Fig 9***.***C)***. In contrast to transcript levels, no change in HIF1A protein was noted for either undifferentiated SHEF1, SHEF2, ZK2012L, or their differentiated progeny. HIF1A protein displayed a slight increase at day 5 differentiation in 2% PG and 2% WKS, and on day 20 in 2% WKS (***Fig 8)***.

**Figure 8.**
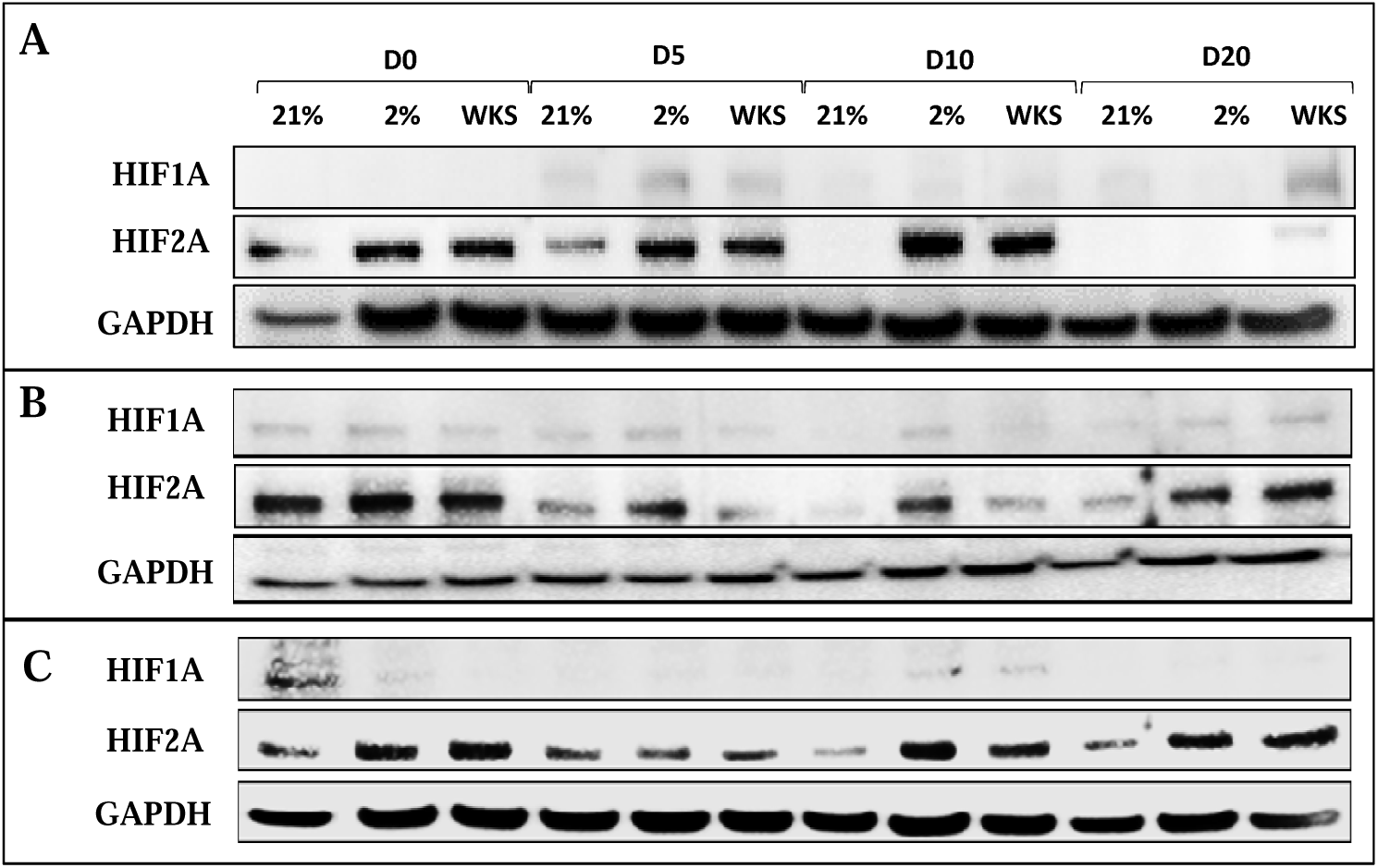
HIF1A and HIF2A protein expression in. **(A)** SHEF1 **(B)** SHEF2 **(C)** ZK2012L. Protein samples were isolated from undifferentiated PSCs and days 5, 10, and 20 differentiated cells exposed to either air oxygen or physoxia conditions. GAPDH is included as a loading control.

**Figure 9.**
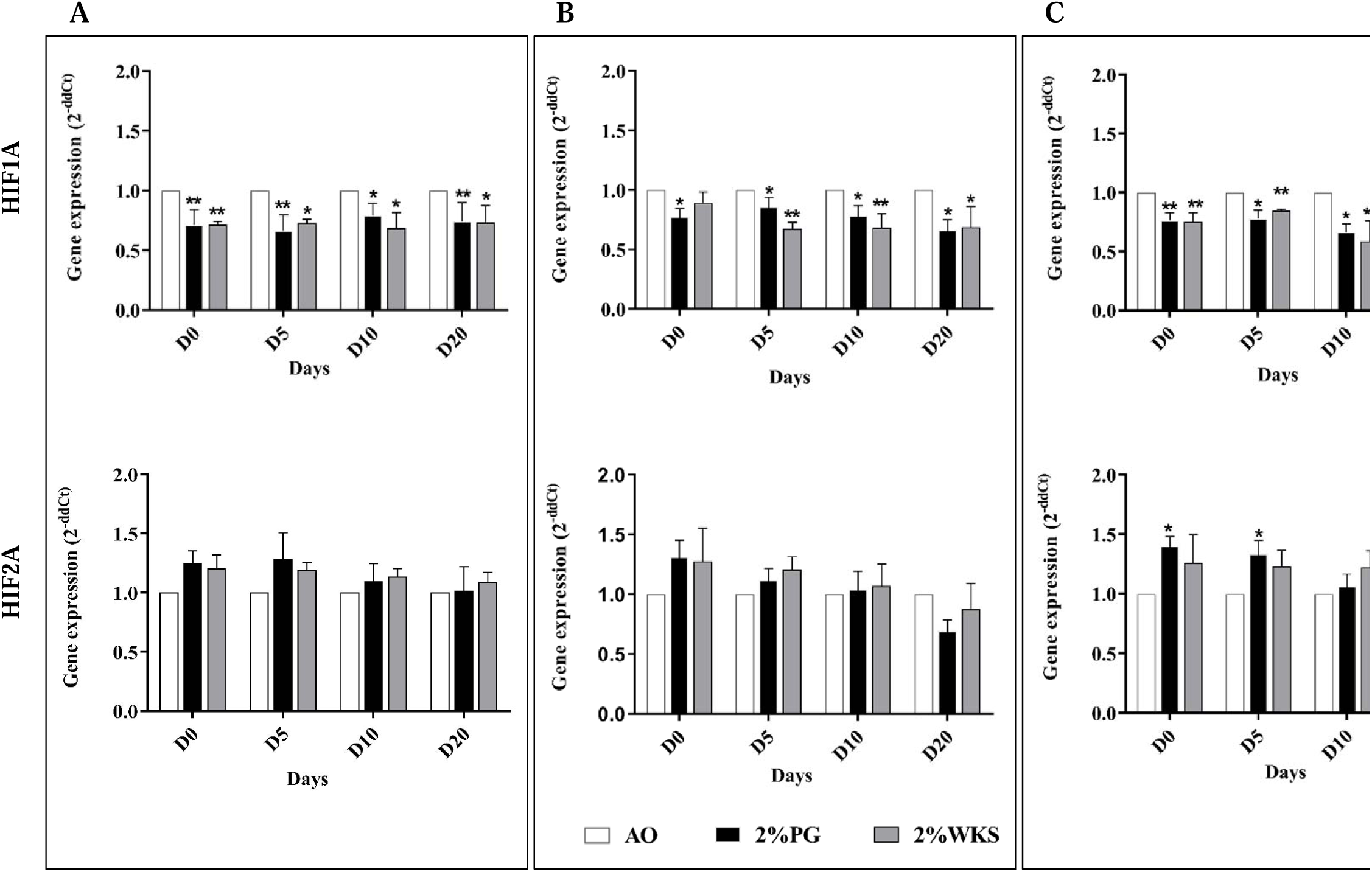
Gene expression of HIF1A and HIF2A in. **(A)** SHEF1, **(B)** SHEF2, and **(C)** ZK2012L. The RT-qPCR expression of the HIFs normalized to the expression of internal control ACTB. Y-axis shows the relative changes in *2*^-ΔΔ*CT*^ of air oxygen exposed cells to physoxia cultured cells. The X-axis indicates time (days). Data are presented as mean (n=3), *P<0.05, **p<0.01 vs 21% AO, error bars indicate standard deviation (SD).

No significant changes to HIF2A gene expression in SHEF1 or SHEF2 were noted. ZK2012L displayed increased expression of HIF2A in undifferentiated and day 5 differentiated cells (p<0.05) in 2% PG only (***Fig 9)***. In contrast, the level of HIF2A protein was increased in undifferentiated SHEF1 and days 5 and 10 differentiated cells in 2%PG and 2%WKS compared to 21% AO. A significant increase in HIF2A protein expression was noted in 2% PG and 2% WKS conditions at day 10 differentiation in SHEF2 cells. Furthermore, there was a significant increase in HIF2A protein level in undifferentiated ZK2012L cells, and in days 10 and 20 differentiation in 2% PG and 2% WKS (***Fig 8)***.

## Discussion

Epigenetics marks including DNA methylation (5mC and 5hmC) impact cellular characteristics, fate, and are critical for various cellular processes ^36^. The role of methylation of Cytosine residues (5mC) at CpG islands is well studied across a range of biological processes including gene expression, cell differentiation, and development ^37^, however, 5hmC is still being explored in cell differentiation and transcriptional regulation in mammalian cells ^38^. Oxygen variation can mediate stem cell fate where, for example, low, physiologic, oxygen conditions increase clonal recovery, maintain a higher number of colony initiating cells, change metabolism and morphology, and enhance proliferation and stemness of PSCs ^22,23,25,39^. Here, we focused on the role of physoxia on PSC characteristics in conjunction with global DNA methylation, the regulation of methylation enzymes, and the methylation of their promoters in human PSCs. We determined, consistent with others, that reduced oxygen conditions enhanced the proliferation and metabolic activity of PSCs when compared to air oxygen cultured cells^19,40^. We observed that global methylation is oxygen-sensitive and associated with the regulation of DNMT3B transcription and translation.

Immunocytochemistry and flow cytometry demonstrated that undifferentiated PSCs expressed the anticipated pluripotency markers; OCT-4, NANOG, ALP, SSEA-4, TRA-1-60, and were negative for SSEA-1. Previous reports suggested that reduced oxygen culture conditions have no detrimental effect on expression of stem cell markers in PSCs ^19,23,41,42^. We noted elevated OCT-4 and SOX-2, not NANOG, expression in 2% WKS condition with flow cytometry determining increased TRA-1-60 expression in reduced oxygen conditions. Therefore, we believe that a controlled oxygen environment 2% WKS provides a more consistent environment for PSCs than 2% PG. Consistent with our observations, hESCs exposed to AO display decreased proliferation and reduced expression of SOX2, NANOG, and OCT4 gene and protein expression when compared with those cultured at 5% O_2_ tri-gas incubators ^19,43^. In contrast to the above, other reports detail no significant difference in SOX2, NANOG, or OCT4 expression in hESCs cultured at 2-4% O_2_ (again with tri-gas incubators) compared to AO ^44,45^.

DNA methylation and hydroxymethylation have essential roles in PSC function and characteristic determination in physiological oxygen conditions ^29,46^. Oxygen can alter the global epigenetic landscape, for instance, global histone methylation was decreased at 5% oxygen tension in hPSCs ^47^. These observations are consistent with ours where we noted decreased global methylation (vs. AO) in 5mC and 5hmC levels in undifferentiated hPSCs exposed to physiological oxygen conditions (2% PG and 2% WKS). Recent reports detail that both 5mC and 5hmC levels are high in ESCs and that these reduce after differentiation ^17,48–51^. Furthermore, essential transcription factors involved in ESCs maintenance of pluripotency such as OCT-4 and NANOG are unmethylated in undifferentiated cells becoming methylated as differentiation progresses ^27,52,53^. We also noted progressively decreased global 5mC and 5hmC levels during differentiation of PSCs and reduced global methylation in physiological oxygen conditions (2% PG and 2% WKS) in differentiated stem cells compared to AO.

Notably, global methylation levels correlated with decreased DNMT3B and TET1 expression in reduced oxygen conditions. ESCs exhibit robust de novo methyltransferase enzyme (DNMT3A and DNMT3B) expression while undifferentiated, with levels of DNMT3B decreasing with progressive differentiation states ^16,54–56^. A previous report has described that expression of DNMT3B was decreased at 5% oxygen in undifferentiated hPSCs ^47^. Consistent with this, we noted reduced DNMT3B expression in physiological oxygen conditions in undifferentiated hPSCs and subsequently differentiated cells. Methylation levels on the DNMT3B promoter were significantly higher under reduced oxygen settings when compared to AO. CpG island methylation can impair transcription factor binding, recruit repressive methyl-binding proteins, and stably silence gene expression ^26^. Taken together, we have evidenced reduced transcriptional expression of DNMT3B in association with increased methylation of its promoter. In contrast, there was a significant decrease in TET1 mRNA level but no significant changes on its promoter CpG content in physoxia versus AO.

Long term exposure of hESC to low oxygen decreased the expression of HIF1A mRNA in hESCs ^19^. We also noted decreased HIF1A gene expression in physoxia. In contrast, long-term physoxic culture resulted in upregulated HIF2A at both transcript and protein levels. Consistent with this observation previous reports have detailed that HIF2A was upregulated following long-term culture of hESC in 5% O_2_ when compared with air cultured counterparts^19^. Further, and indicative of a conserved behaviour, HIF2A was upregulated at transcriptional and translational levels in human MSCs in a 5% O_2_ culture setting ^57^. Repression of HIF2A via siRNA resulted in decreased proliferation and reduced expression of OCT-4, NANOG and SOX-2 in hESCs cultured under 5% O_2_ ^19^. In addition, DNMT1 and DNMT3B promoters each display hypoxia response elements for HIF-binding ^33,46^. Therefore, we hypothesise that increased HIF2A and reduced HIF1A under physoxia may associate with, or even drive, the observed decrease in DNMT3B overall the increase in its promoter methylation.

In summary, we demonstrate that physiological oxygen tension increased pluripotency marker expression, proliferation, and metabolic activity with a concurrent decreased DNMT3B, TET1 gene expression, and global DNA methylation in hPSCs. Further, DNMT3B expression was methylation-regulated in an oxygen-sensitive manner. We show that physoxia conditions modify levels of global methylation linked to DNMT3B expression and that hPSCs display oxygen-sensitive methylation patterns. Importantly, air oxygen is almost never normoxic in mammalian cell culture and increased global methylation can be considered as an artefact of air oxygen culture. Physoxia is an essential component for regenerative medicine to safely realise cell potential and enable clean characterisation.

## Abbreviations

(5mC): 5-Methylcytosine and
(5hmC): 5-hydroxymethylcytosine
(PSCs): pluripotent stem cells
(ESCs): embryonic stem cells
(MSCs): mesenchymal stem cells
(HIF)-1: hypoxia-inducible factor
(BM): bone marrow
(DNMT): DNA Methyltransferase
(HDAC): Histone deacetylases
(TET) enzymes: Ten eleven translocation

## Acknowledgements

The work described in this paper was principally funded by support from the Iraqi Ministry of Higher Education and Scientific Research, University of Baghdad, Baghdad, Iraq (S1453). We also wish to thank the Turkish Ministry of National Education for their support. Induced pluripotency cell line ZK2012L was kindly provided by Prof Susan Kimber.

