## Supplementary figures and images for "Physoxia influences global and gene-specific methylation in pluripotent stem cells"

### Supplementary Figure 1

#### Slide 1
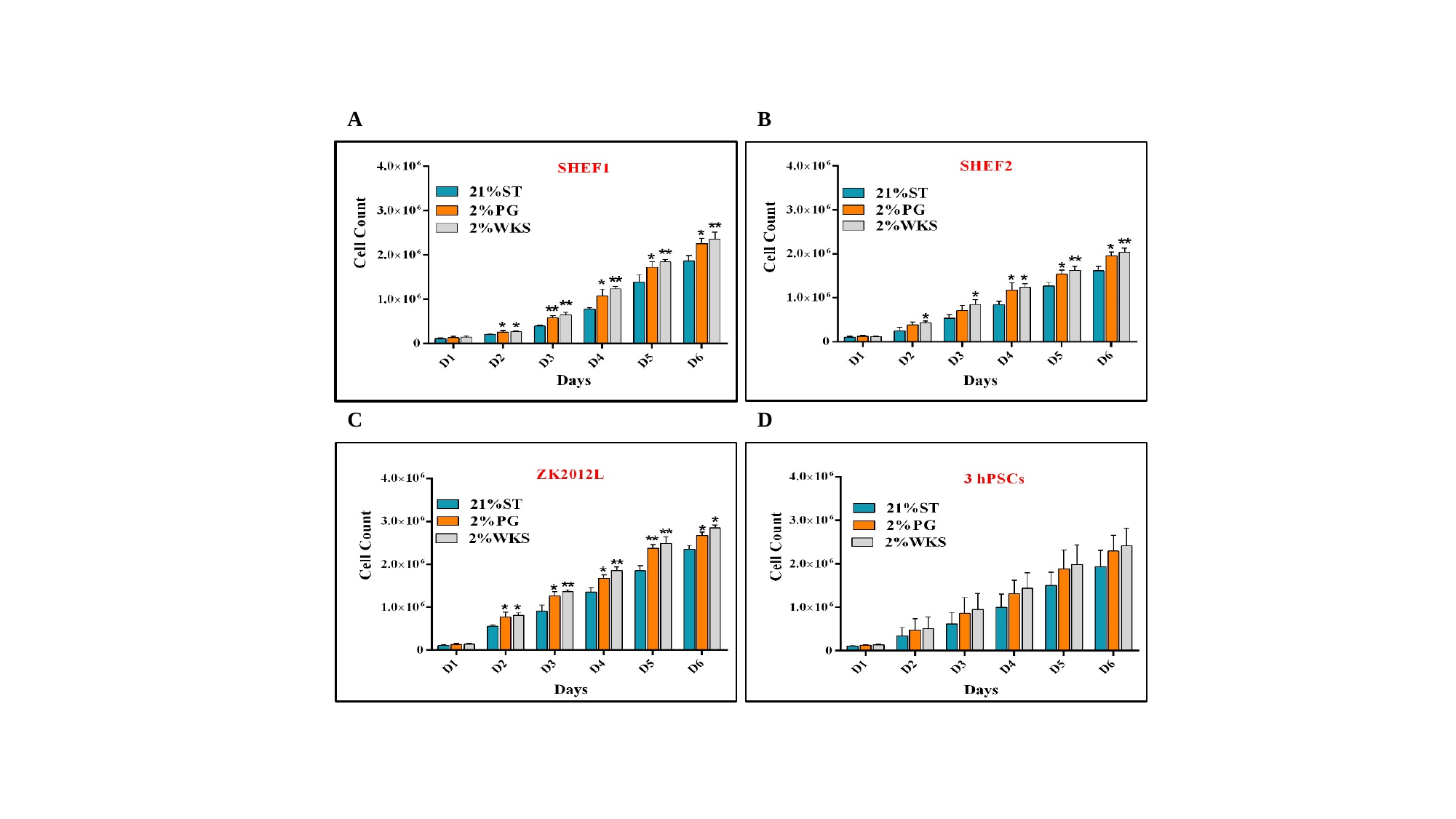

B
A
C
D

#### Slide 2
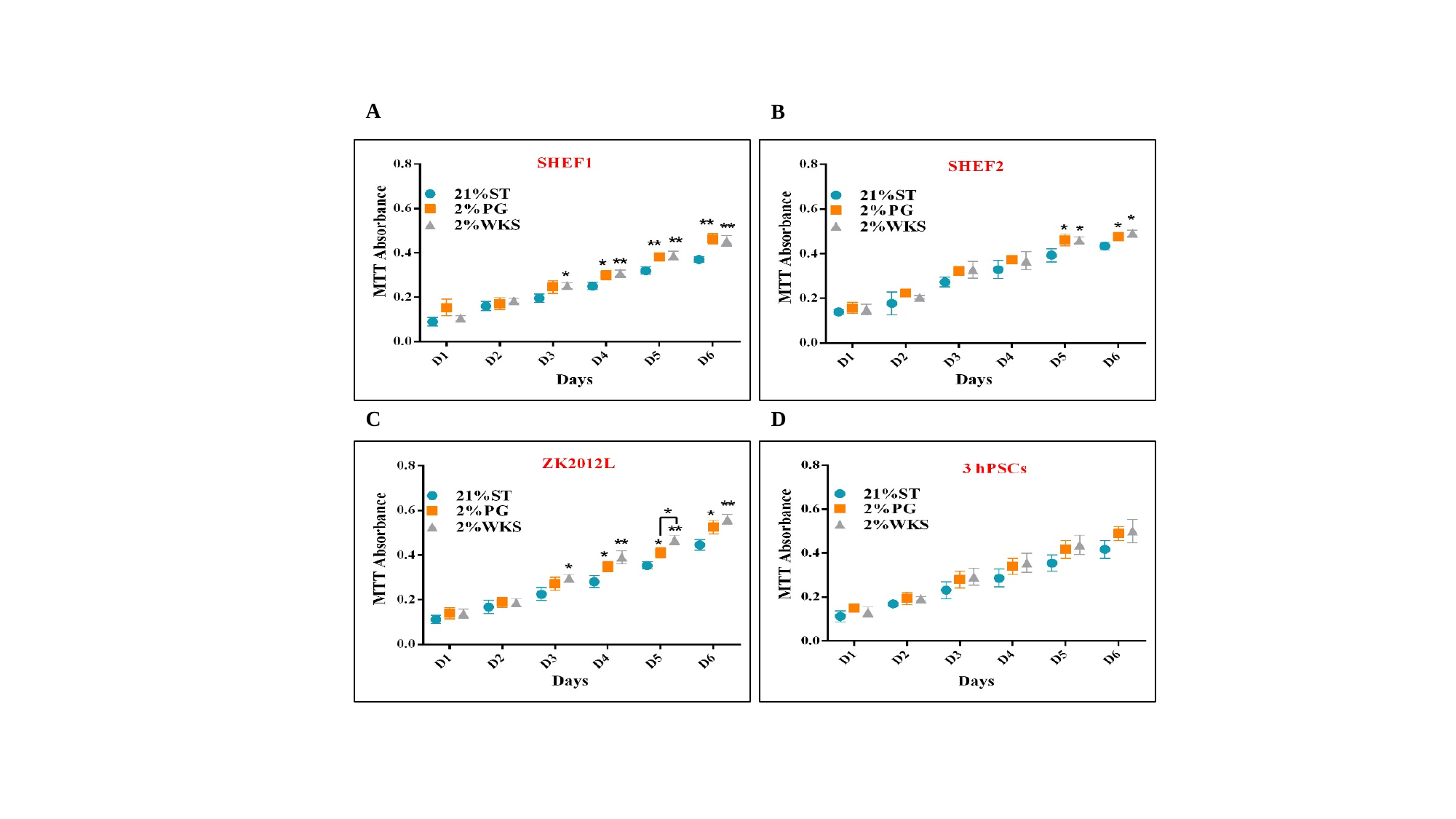

A
B
C
D

#### Slide 3
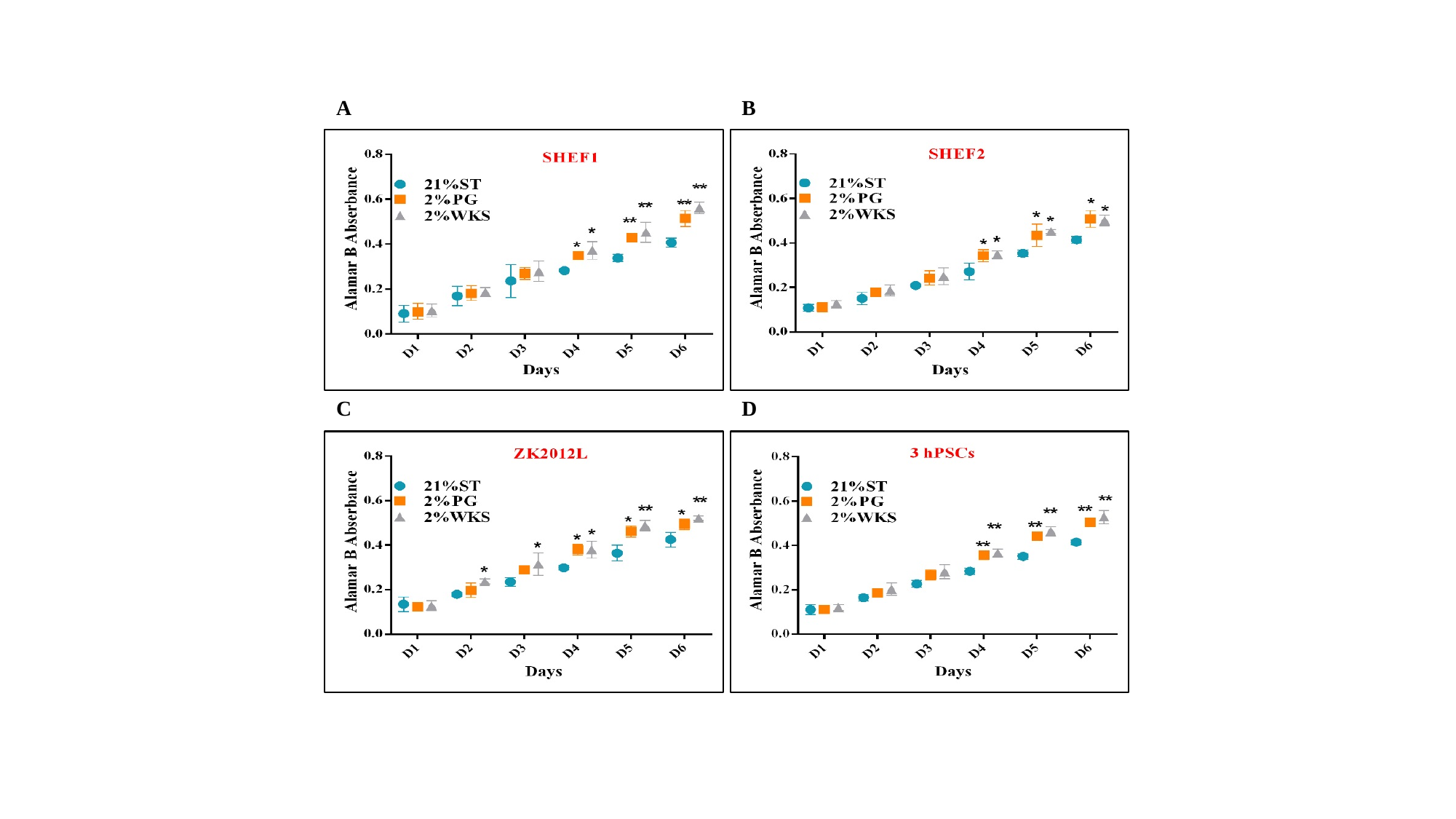

A
B
C
D
