## Supplementary Figure 2 for "Physoxia influences global and gene-specific methylation in pluripotent stem cells"

### Slide 1
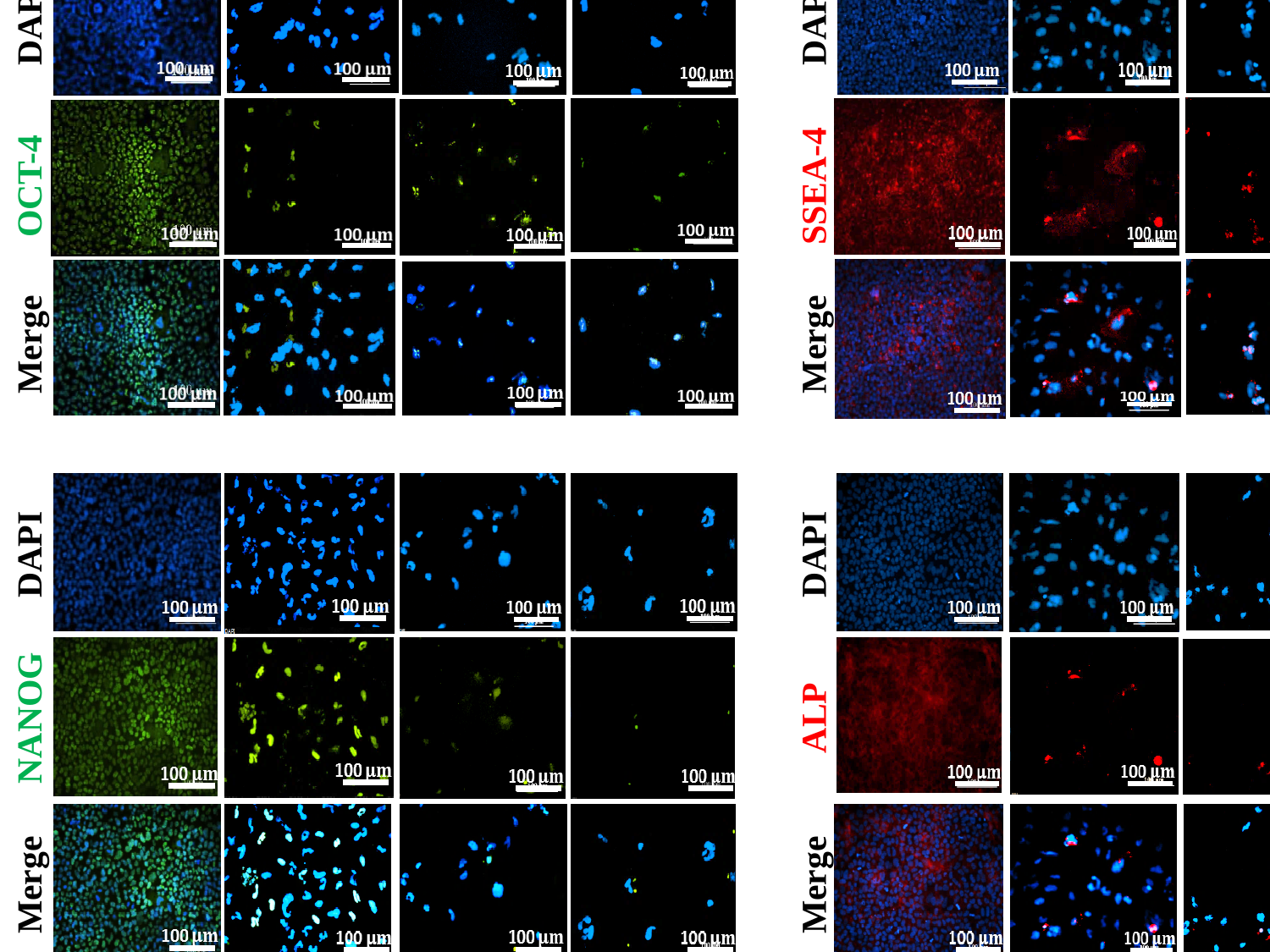

SHEF1, 21%ST
DAY 0
DAY 5
DAY 10
DAY 20
DAY 0
DAY 5
DAY 10
DAY 20
DAPI
DAPI
OCT-4
SSEA-4
Merge
Merge
DAPI
DAPI
NANOG
ALP
Merge
Merge

### Slide 2
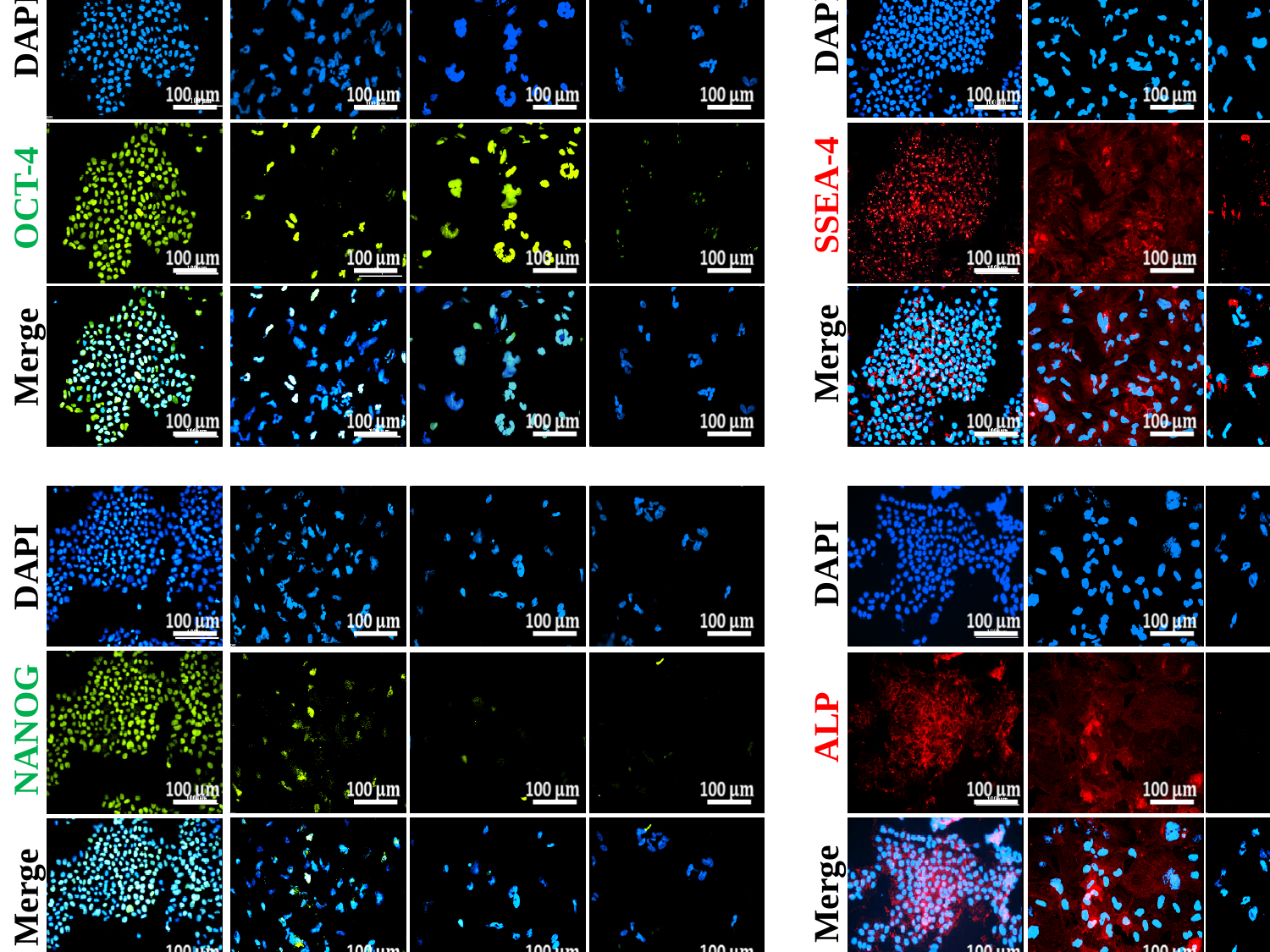

SHEF1, 2%PG
DAY 0
DAY 5
DAY 10
DAY 20
DAY 0
DAY 5
DAY 10
DAY 20
DAPI
DAPI
SSEA-4
OCT-4
Merge
Merge
DAPI
DAPI
ALP
NANOG
Merge
Merge

### Slide 3
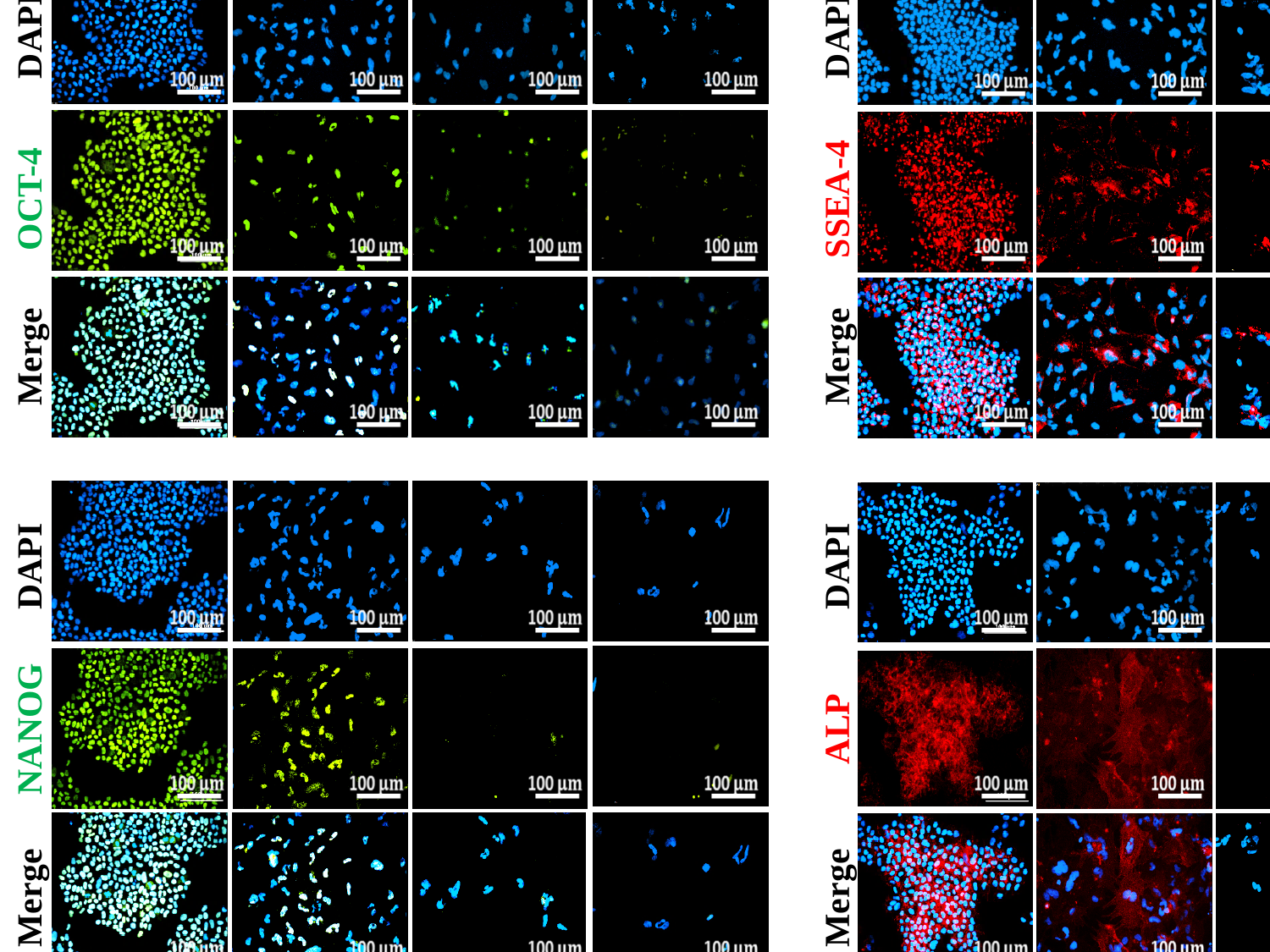

SHEF1, 2%WKS
DAY 0
DAY 5
DAY 10
DAY 20
DAY 0
DAY 5
DAY 10
DAY 20
DAPI
DAPI
OCT-4
SSEA-4
Merge
Merge
DAPI
DAPI
NANOG
ALP
Merge
Merge

### Slide 4
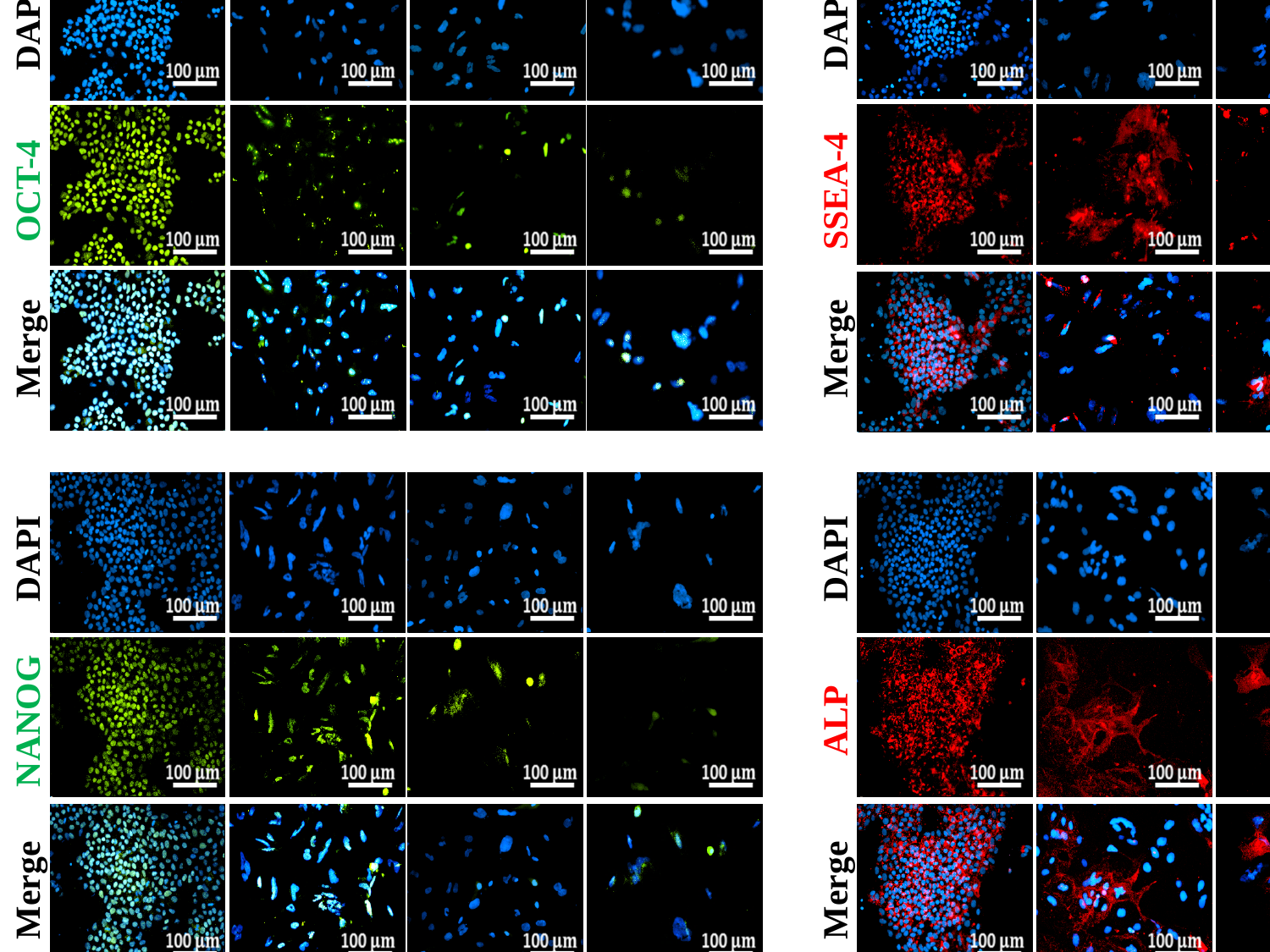

SHEF2, 21%ST
DAY 0
DAY 5
DAY 10
DAY 20
DAY 0
DAY 5
DAY 10
DAY 20
DAPI
DAPI
OCT-4
SSEA-4
Merge
Merge
DAPI
DAPI
NANOG
ALP
Merge
Merge

### Slide 5
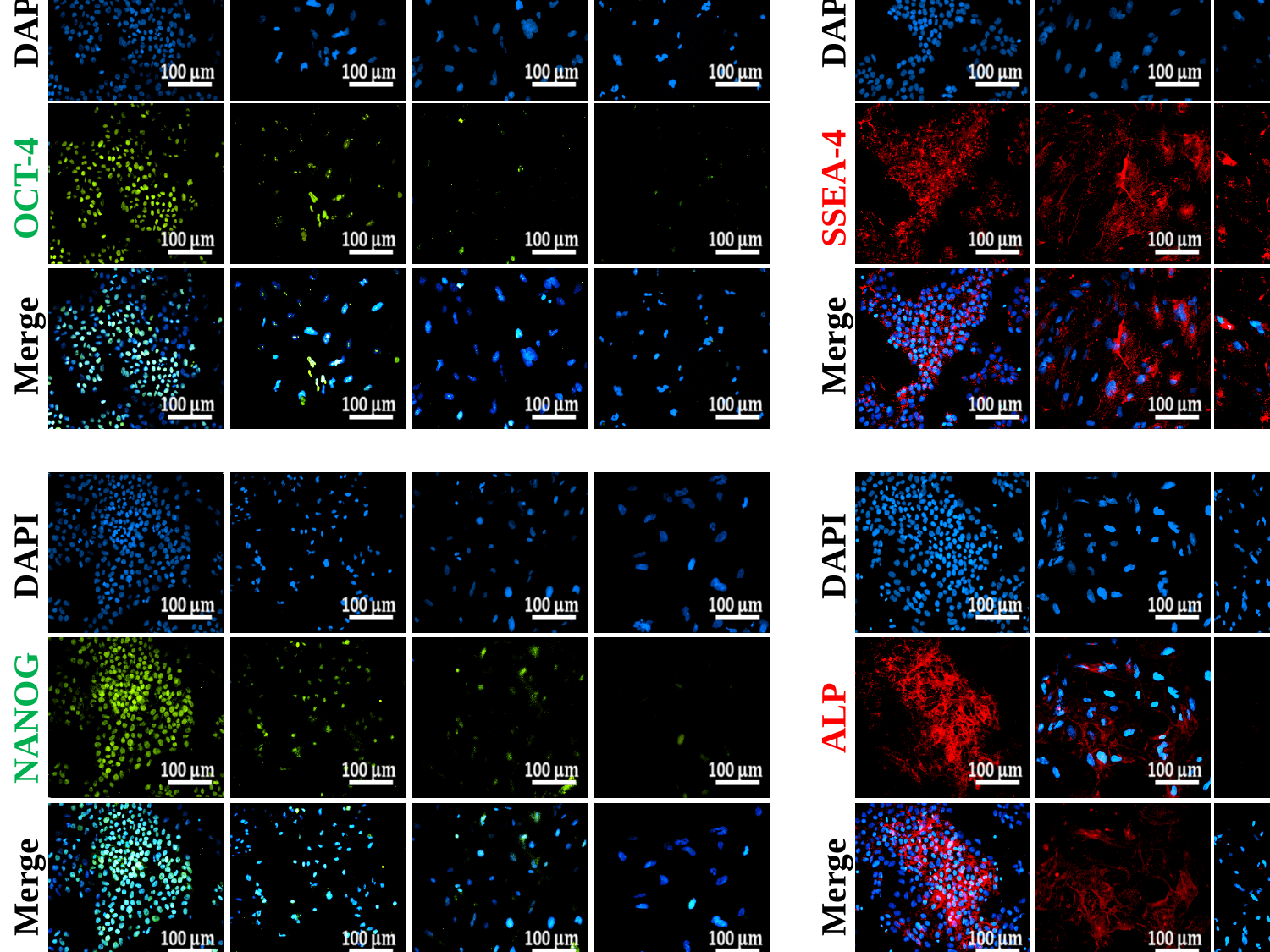

SHEF2, 2%PG
DAY 0
DAY 5
DAY 10
DAY 20
DAY 0
DAY 5
DAY 10
DAY 20
DAPI
DAPI
OCT-4
SSEA-4
Merge
Merge
DAPI
DAPI
NANOG
ALP
Merge
Merge

### Slide 6
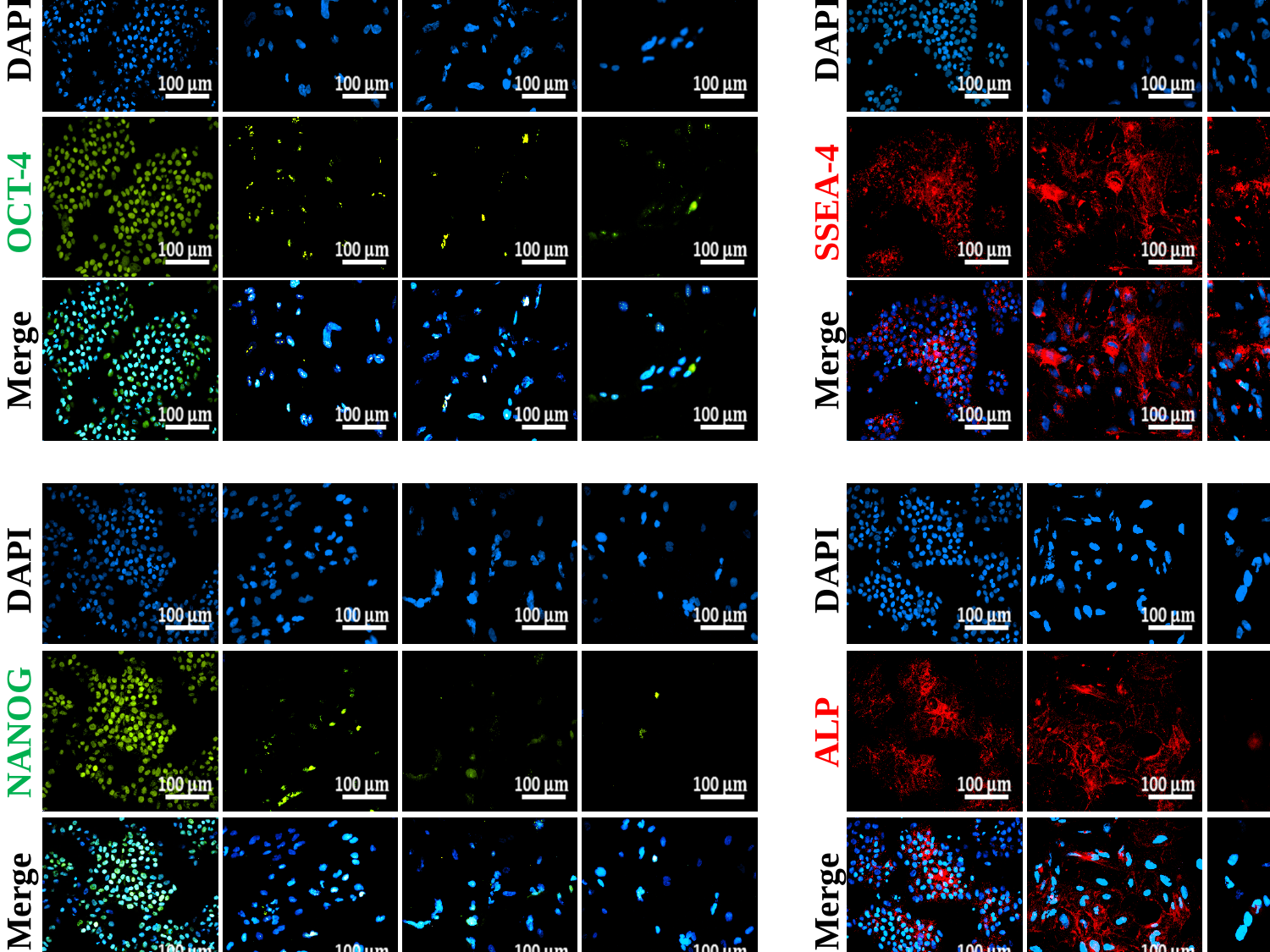

SHEF2, 2%WKS
DAY 0
DAY 5
DAY 10
DAY 20
DAY 0
DAY 5
DAY 10
DAY 20
DAPI
DAPI
OCT-4
SSEA-4
Merge
Merge
DAPI
DAPI
NANOG
ALP
Merge
Merge

### Slide 7
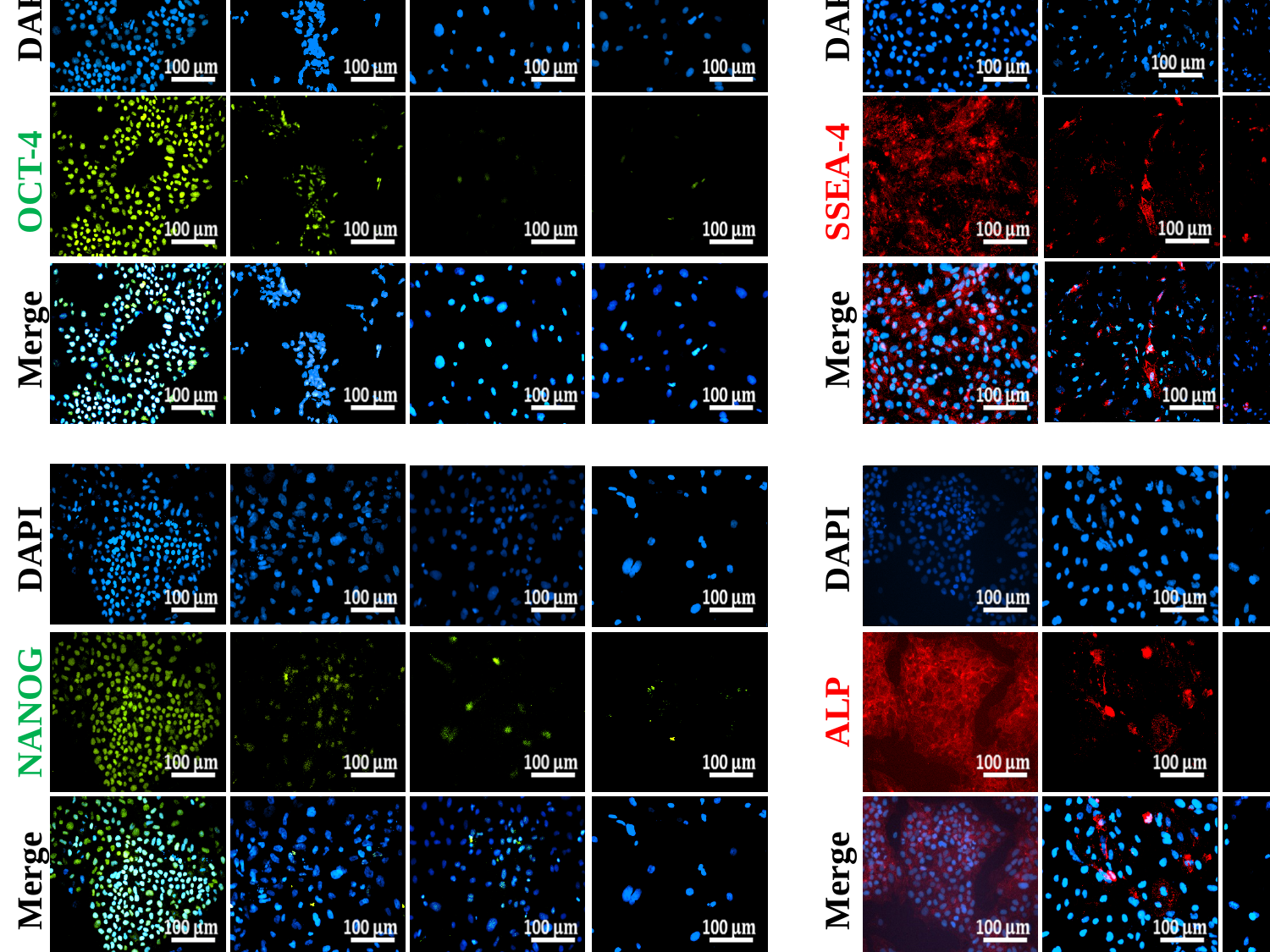

DAY 0
DAY 5
DAY 10
DAY 20
DAY 0
DAY 5
DAY 10
DAY 20
DAPI
DAPI
OCT-4
SSEA-4
Merge
Merge
DAPI
DAPI
NANOG
ALP
Merge
Merge
IPSCs, 21%ST

### Slide 8
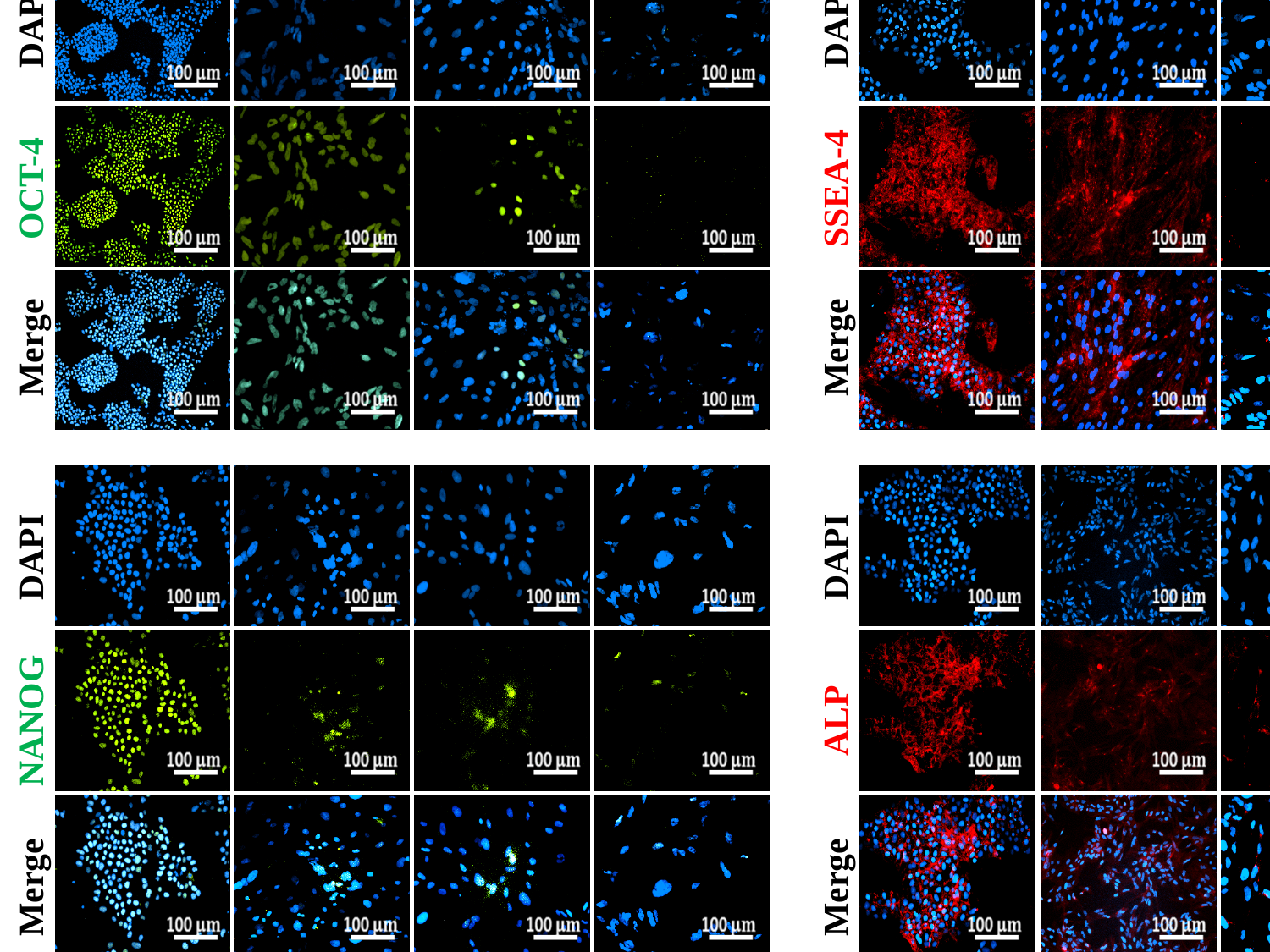

IPSCs, 2%PG
DAY 0
DAY 5
DAY 10
DAY 20
DAY 0
DAY 5
DAY 10
DAY 20
DAPI
DAPI
OCT-4
SSEA-4
Merge
Merge
DAPI
DAPI
NANOG
ALP
Merge
Merge

### Slide 9
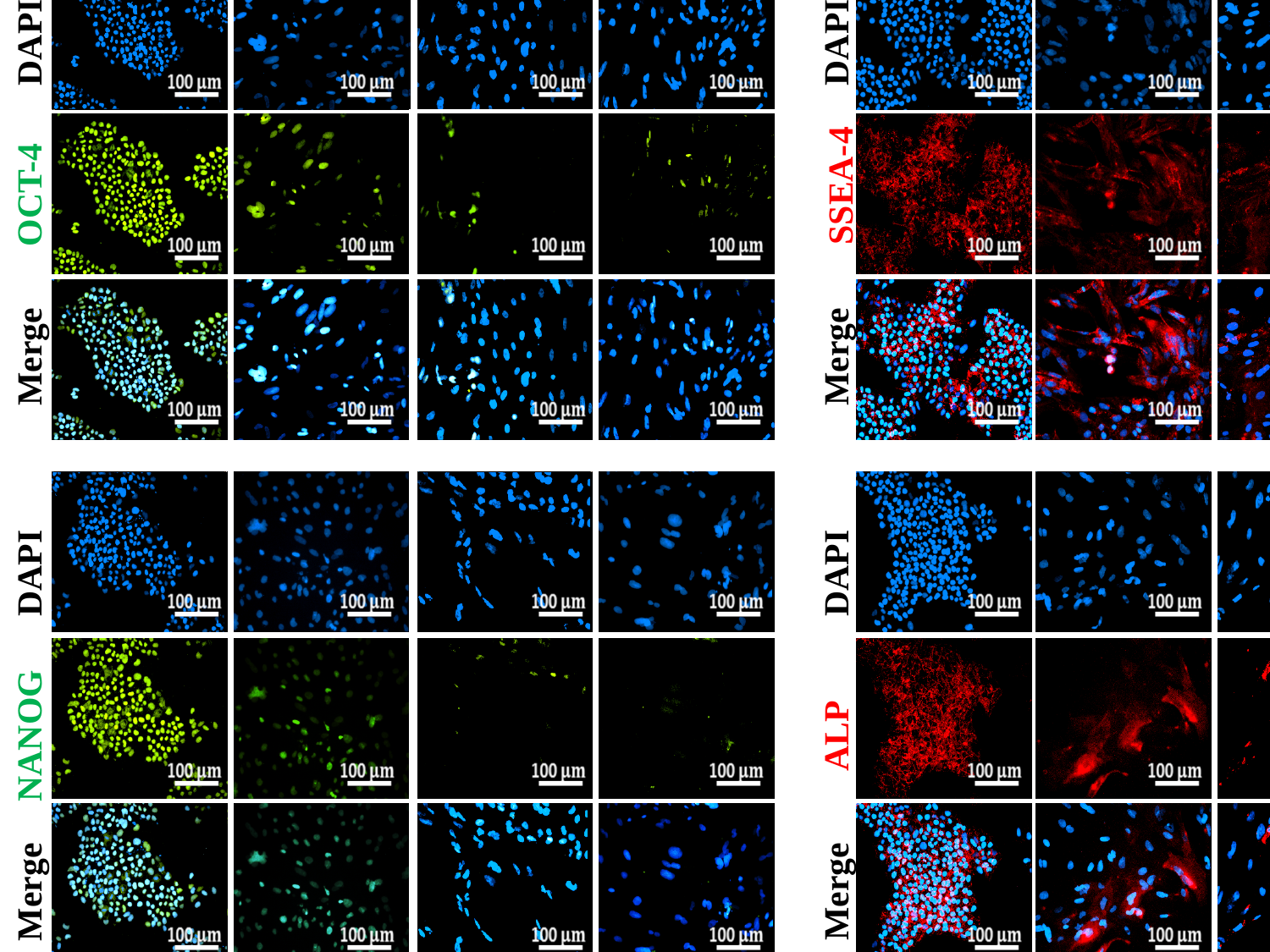

IPSCs, 2%WKS
DAY 0
DAY 5
DAY 10
DAY 20
DAY 0
DAY 5
DAY 10
DAY 20
DAPI
DAPI
SSEA-4
OCT-4
Merge
Merge
DAPI
DAPI
NANOG
ALP
Merge
Merge

### Slide 10
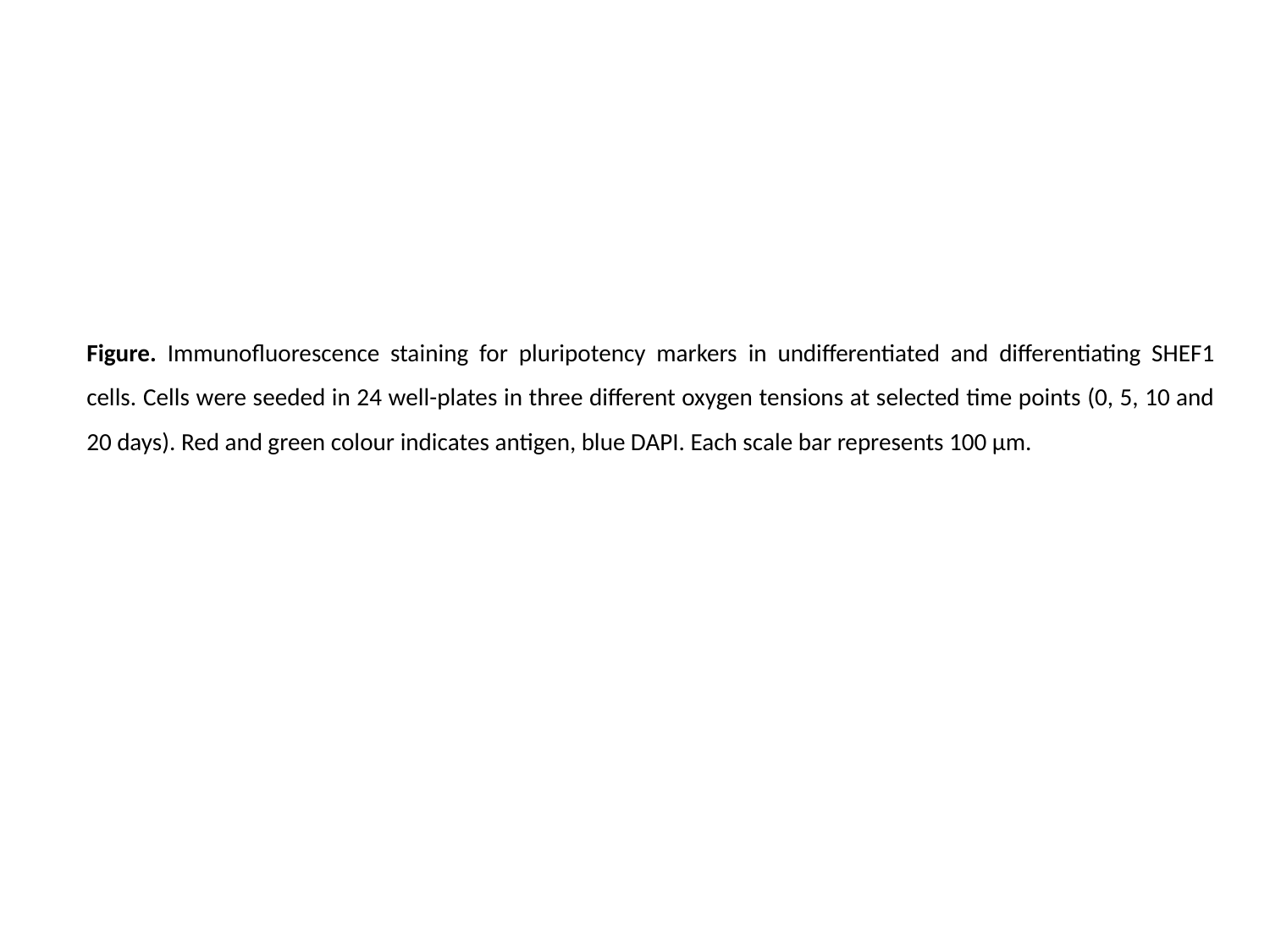

Figure. Immunofluorescence staining for pluripotency markers in undifferentiated and differentiating SHEF1 cells. Cells were seeded in 24 well-plates in three different oxygen tensions at selected time points (0, 5, 10 and 20 days). Red and green colour indicates antigen, blue DAPI. Each scale bar represents 100 µm.
